# MGDrivE 2: A simulation framework for gene drive systems incorporating seasonality and epidemiological dynamics

**DOI:** 10.1101/2020.10.16.343376

**Authors:** Sean L. Wu, Jared B. Bennett, Héctor M. Sánchez C., Andrew J. Dolgert, Tomás M. León, John M. Marshall

**Author notes:** **Correspondence:** Sean L. Wu John M. Marshall.

## Abstract

1. Interest in gene drive technology has continued to grow as promising new drive systems have been developed in the lab and discussions are moving towards implementing field trials. The prospect of field trials requires models that incorporate a significant degree of ecological detail, including parameters that change over time in response to environmental data such as temperature and rainfall, leading to seasonal patterns in mosquito population density. Epidemiological outcomes are also of growing importance, as: i) the suitability of a gene drive construct for release will depend on its expected impact on disease transmission, and ii) initial field trials are expected to have a measured entomological outcome and a modeled epidemiological outcome.
2. We present MGDrivE 2 (Mosquito Gene Drive Explorer 2): an extension of and development from the MGDrivE 1 simulation framework that investigates the population dynamics of a variety of gene drive architectures and their spread through spatially-explicit mosquito populations. Key strengths and improvements of the MGDrivE 2 framework are: i) the ability of parameters to vary with time and induce seasonal population dynamics, ii) an epidemiological module accommodating reciprocal pathogen transmission between humans and mosquitoes, and iii) an implementation framework based on stochastic Petri nets that enables efficient model formulation and flexible implementation.
3. Example MGDrivE 2 simulations are presented to demonstrate the application of the framework to a CRISPR-based homing gene drive system intended to drive a disease-refractory gene into a population, incorporating time-varying temperature and rainfall data, and predict impact on human disease incidence and prevalence. Further documentation and use examples are provided in vignettes at the project’s CRAN repository.
4. MGDrivE 2 is an open-source R package freely available on CRAN. We intend the package to provide a flexible tool capable of modeling gene drive constructs as they move closer to field application and to infer their expected impact on disease transmission.

## 1. Introduction

Interest in gene drive technology has continued to grow in recent years as a range of promising new constructs have been developed in the lab and discussions have moved towards implementing field trials in some cases. Recently developed systems include a CRISPR-based homing system intended for population suppression targeting the *doublesex* gene in *Anopheles gambiae*, the main African malaria vector (Kyrou *et al.*, 2018), a split gene drive system intended for confineable and transient population replacement in *Aedes aegypti*, the main vector of dengue, chikungunya and Zika viruses (Li *et al.*, 2020), and CRISPR-based homing systems intended for population replacement in *An. gambiae* (Carballar-Lejarazú *et al.*, 2020) and *Anopheles stephensi*, the main malaria vector in urban India (Adolfi *et al.*, 2020).

As the technology advances and potential field trials are discussed (James *et al.*, 2018), models are needed that incorporate additional ecological detail, including parameters that change over time in response to environmental variables such as temperature and rainfall, as well as models linking entomological and epidemiological outcomes (James *et al.*, 2020). Many insects, including mosquitoes, display a high degree of seasonality in their population dynamics, as development time from one life stage to another, and mortality rates associated with each life stage, vary with temperature and other environmental variables (Mordecai *et al.*, 2019). For *An. gambiae* and several other mosquito disease vectors, population size varies largely in response to recent rainfall, which creates pools of standing water and hence enhanced carrying capacity of the environment for mosquito larvae (White *et al.*, 2011). Seasonal changes in temperature and rainfall thus lead to seasonal changes in mosquito population density and consequent disease transmission, which must be accounted for in disease control strategies.

Models of disease transmission are also becoming increasingly relevant to models of gene drive dynamics, as: i) the readiness of a gene drive system for field trials will be determined in part by its expected (i.e. modeled) epidemiological impact, and ii) initial field trials are expected to have a measured entomological outcome alongside a modeled epidemiological outcome (James *et al.*, 2018). Given the potential for a non-localized gene drive system to spread broadly, it has been acknowledged that constructs at the trial stage should be expected to cause a significant reduction in disease transmission, as even a confined trial could lead to wide-scale spread for an effective system (James *et al.*, 2018). Therefore, readiness for field trials should be determined by alignment with a target product profile (TPP) and/or list of preferred product characteristics (PPCs) that include expected impact on disease transmission (James *et al.*, 2020). Models that incorporate both gene drive and epidemiological dynamics can account for local malaria or arboviral transmission dynamics and specify gene drive construct parameters that achieve the desired level of epidemiological control.

Previously, we developed the MGDrivE 1 modeling framework to model the population dynamics of a variety of genetics-based and biological control systems and their spread through spatially-explicit populations of mosquitoes, or insects having a similar life history (Sánchez *et al.*, 2020). Here, we present MGDrivE 2, which significantly improves upon the capabilities of MGDrivE 1 by addressing the above-mentioned considerations, namely: i) the ability of parameter values to change over time, and hence to model seasonal population dynamics, and ii) the incorporation of an epidemiology module that can accommodate pathogen transmission between humans and mosquitoes. Minor additional improvements have been made to the inheritance, life history and landscape modules of the framework to reflect advances in these fields; for instance, a more resolved understanding of maternal deposition of Cas protein for CRISPR-based gene drive systems has been incorporated (Champer *et al.*, 2018). Models in MGDrivE 2 are represented as a stochastic Petri net (SPN), which has both computational and architectural benefits: model specification is separate from simulation, models can be efficiently stored and updated in memory, and a wealth of fast simulation algorithms from other fields can be used (Goss and Peccoud, 1998).

In this paper, we describe the key developments implemented in MGDrivE 2. We then demonstrate the application of the framework to the disease control impact of a CRISPR-based homing gene drive system intended to drive a disease-refractory gene into a population, and conclude with a discussion of future needs and applications for simulation packages in the field of gene drive modeling.

## 2. Design and implementation

MGDrivE 2 is a significant extension of and development from MGDrivE 1, a model for the spread of gene drive systems through spatially-explicit mosquito populations. The MGDrivE 2 model incorporates: i) an “inheritance module” that describes the distribution of offspring genotypes for given maternal and paternal genotypes, ii) a “life history module” that describes the development of mosquitoes from egg to larva to pupa to adult, iii) a “landscape module” that describes the distribution and movement of mosquitoes through a metapopulation, and iv) an “epidemiology module” that describes pathogen transmission between mosquitoes and humans (Figure 1). The framework is formulated as a SPN that can be mapped to a continuous-time Markov process in which model parameters may vary over time. It can also be implemented as a deterministic model via mean-field approximation of the stochastic model (Bortolussi *et al.*, 2013).

**Figure 1.**
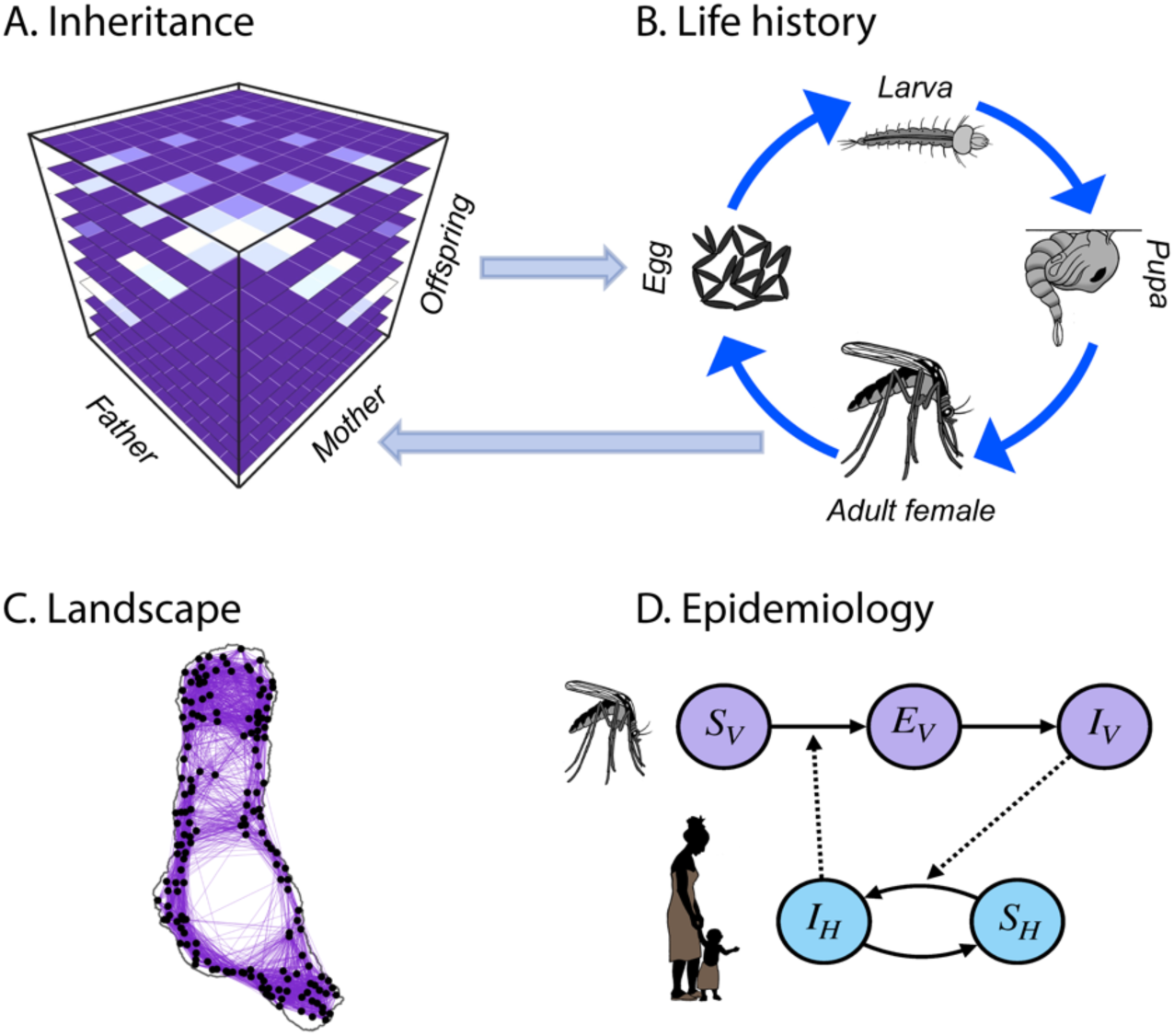
Modules in the MGDrivE 2 framework. **(A)** Genetic inheritance is embodied by a three-dimensional tensor referred to as an “inheritance cube.” Maternal and paternal genotypes are depicted on the *x* and *y*-axes and offspring genotypes on the *z*-axis. **(B)** Mosquito life history is modeled according to an egg-larva-pupa-adult (female and male) life cycle in which density dependence occurs at the larval stage, and life cycle parameters may vary as a function of environmental variables over time. Genotypes are tracked across all life stages, and females obtain a composite genotype upon mating - their own and that of the male they mate with. Egg genotypes are determined by the inheritance cube. **(C)** The landscape represents a metapopulation in which mosquitoes are distributed across population nodes and move between them according to a dispersal kernel. Population sizes and movement rates may vary as a function of environmental variables. **(D)** The epidemiology module describes reciprocal transmission of a vector-borne pathogen between mosquitoes and humans. This requires modeling human as well as mosquito populations, and the number of individuals having each infectious state. Epidemiological parameters may vary as a function of environmental variables.

The core framework is developed in R (https://www.r-project.org/). The SPN framework enables separation of model components, allowing users to modify code on a component-by-component basis as needed for model specification or computational speed. We now describe the model extensions and developments from MGDrivE 1 to 2 in more detail. Full details of the MGDrivE 2 model framework are provided in the S1 Text.

### 2.1. Time-dependent parameters and seasonality

The incorporation of time-dependent parameters represents a significant improvement of the MGDrivE 2 modeling framework. In MGDrivE 1, the mosquito life history module follows the lumped age-class model of Hancock and Godfray as adapted by Deredec *et al.* (2011), which describes development from egg to larva to pupa to adult using delay-difference equations. The delay framework allows development times to be modeled as fixed rather than exponentially-distributed; however, it is not compatible with time-varying parameters as these could vary during the delay. In MGDrivE 2, the discrete-time, fixed-delay framework of MGDrivE 1 is replaced by a continuous-time implementation in which each life stage is divided into a series of substages. For a single substage, the development time is exponentially-distributed; but as the number of substages increases, the distribution of development times becomes concentrated around the mean. Specifically, if a life stage with a mean development time of 1/*d* is divided into a series of *n* substages, the new development times are Erlang-distributed with mean, *1/d*, and variance, 1/(*dn*2), or equivalently, with shape parameter, *n*, and rate parameter, *d*/*n*. The development time, *d*(*t*), may also vary over time, *t*; however the number of substages, *n*, and hence the mean-variance relationship for development times, must remain constant within a simulation.

Most importantly, the new model implementation allows any model parameter to vary with time, enabling the framework to account for seasonal variation in development times and mortality rates due to environmental dependencies. Temperature is known to strongly influence development times for juvenile mosquito stages, and mortality rates for all mosquito life stages (Beck-Johnson *et al.*, 2017; Mordecai *et al.*, 2019), and rainfall is known to influence the carrying capacity of the environment for larvae, and therefore density-dependent larval mortality rates (White *et al.*, 2011; Muriu *et al.*, 2013). The new model formulation allows these parameters to vary in continuous time in response to environmental data, and hence for seasonal variations in temperature and rainfall to drive seasonal variations in mosquito population density.

Parameters defining other modules of the model - inheritance, landscape and epidemiology - are also able to vary over time within the new model formulation. For instance, gene drive systems under the control of temperature-dependent promoters (Zeidler *et al.*, 2004; Dissmeyer, 2017) may have time-varying homing efficiencies, mosquito movement rates may vary seasonally in response to temperature and other environmental factors (Le Goff *et al.*, 2019), and epidemiological parameters such as the extrinsic incubation period (EIP) and pathogen transmission probabilities from human-to-mosquito and mosquito-to-human are all known to display seasonal variation through temperature dependence (Beck-Johnson *et al.*, 2017; Mordecai *et al.*, 2019).

### 2.2. Epidemiology module

The epidemiology module describes reciprocal transmission of a vector-borne pathogen between mosquitoes and humans. This requires modeling of both vector and human populations, as well as an attribute describing the number of individuals in the vector and human populations having each infectious state (Figure 2). To model malaria, the Ross-Macdonald model is included, which has susceptible (S_V_), exposed/latently infected (E_V_), and infectious (I_V_) states for mosquitoes, and susceptible (S_H_), and infected/infectious (I_H_) states for humans (Ross, 1910; Macdonald, 1957). Malaria infection in humans is described by an SIS model, in which humans become infected at a per-capita rate equal to the “force of infection” in humans, *λ*_*H*_, and recover at a rate, *r*. Malaria infection in mosquitoes is described by an SEI model, in which adult mosquitoes emerge from pupae in the susceptible state, become exposed and latently infected at a per-capita rate equal to the force of infection in mosquitoes, *λ*_*V*_, and progress to infectiousness at a rate equal to *γ*_*V*_. The force of infection in humans, *λ*_*H*_, is proportional to the fraction of mosquitoes that are infectious, *I_V_*/*N_V_*, where *N_V_* is the adult mosquito population size, and the force of infection in mosquitoes, *λ_V_*, is proportional to the fraction of humans that are infectious, *I_H_*/*N_H_*, where *N_H_* is the human population size. Since an exponentially-distributed EIP leads to some mosquitoes having unrealistically brief incubation periods, we divide the EV state into a series of *n* sub-states, as described in section 2.1, leading to the EIP being Erlang-distributed with shape parameter, *n*, and rate parameter, *γ_V_*/*n* (Smith, Dushoff and Ellis McKenzie, 2004). Finally, transmission parameters may be tied to specific mosquito genotypes - for instance, an antimalarial effector gene may be associated with a human-to-mosquito or mosquito-to-human transmission probability of zero.

**Figure 2.**
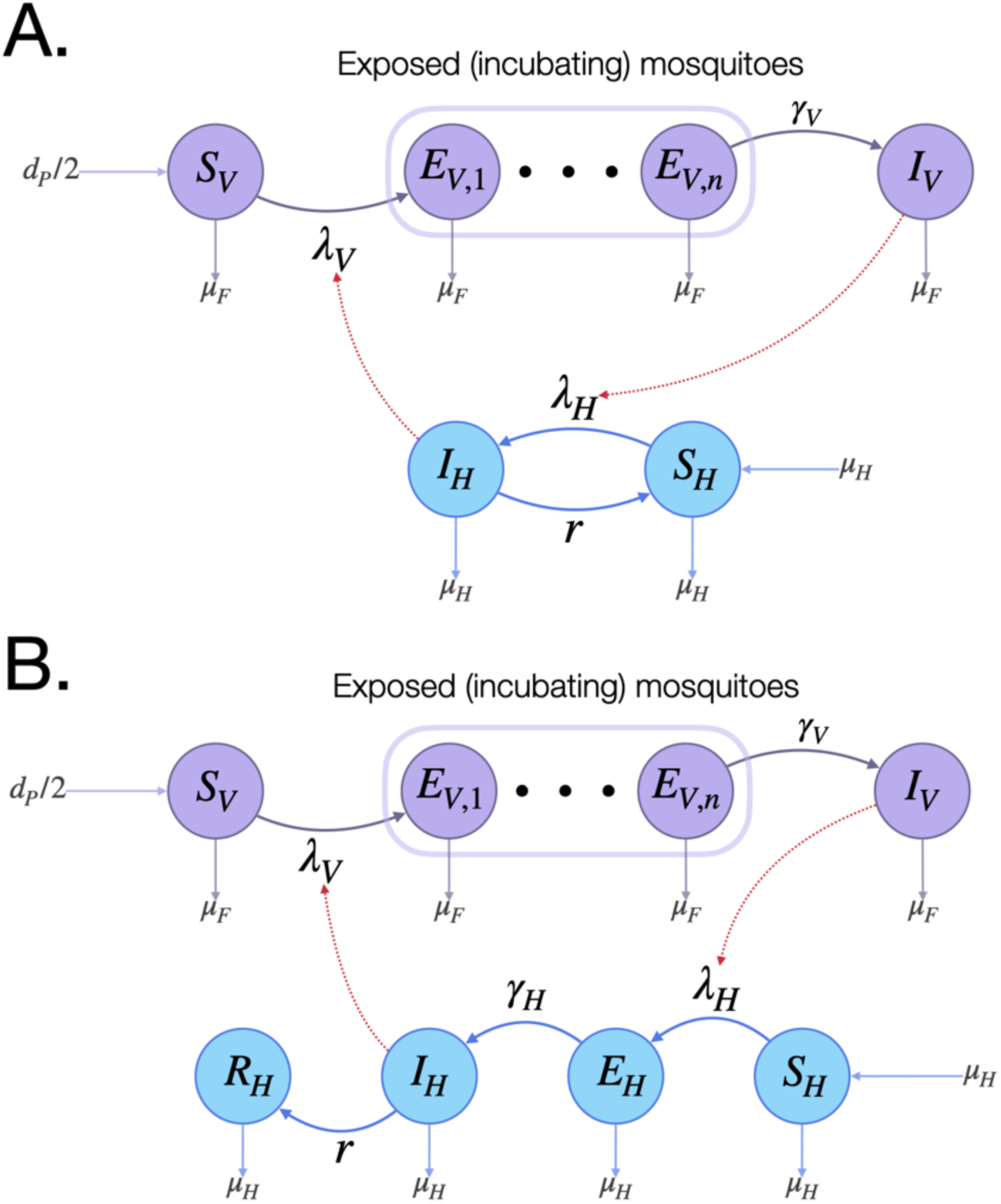
Epidemiology module. MGDrivE 2 includes two basic models for reciprocal pathogen transmission between mosquitoes and humans - one for malaria **(A)**, and one for arboviruses **(B)**. In both cases, female mosquitoes emerge from pupae at a rate equal to *d*_*P*_/2 as susceptible adults (S_V_), become exposed/latently infected (E_V,1_) at a rate equal to the force of infection in mosquitoes, *λ*_*V*_, and progress to infectiousness (I_V_) through the extrinsic incubation period (EIP = 1/*γ*_*V*_), which is divided into *n* bins to give an Erlang-distributed dwell time. The mortality rate, *µ*_*F*_, is the same for female mosquitoes in each of these states. For malaria **(A)**, susceptible humans (S_H_) become infected/infectious (I_H_) at a rate equal to the force of infection in humans, *λ*_*H*_, and recover at rate *r*, becoming susceptible again. For arboviruses **(B)**, susceptible humans (S_H_) become exposed/latently infected (E_H_) at a rate equal to *λ*_*H*_, progress to infectiousness (I_H_) at rate equal to *f*, and recover (R_H_) at rate, *r*. Infection dynamics couple the mosquito and human systems via the force of infection terms; *λ*_*V*_ is a function of I_H_, and *λ*_*H*_ is a function of I_V_, shown via red edges.

To model arboviruses such as chikungunya, Zika and single serotypes of dengue virus, we include an SEIR model for human transmission, in which the human states are: susceptible (S_H_), exposed/latently infected (E_H_), infectious (I_H_), and removed/recovered (R_H_) (Kucharski *et al.*, no date; Robinson *et al.*, 2014). The E_H_ and R_H_ states are included because arboviruses are generally thought to be immunizing, and have latent periods that tend to be on a similar timescale to the duration of infectiousness. Humans become exposed/latently infected at a per-capita rate equal to *λ*_*H*_, progress to infectiousness at rate, *γ*_*H*_, and recover at rate, *r*. For mosquito transmission, the SEI model with an Erlang-distributed EIP is used again. Further details of the mathematical formulation of both the malaria and arbovirus models are provided in the S1 Text. The extensibility of the SPN framework means that more complex epidemiological models can be developed and implemented by users.

Modeling vector-borne disease transmission within a metapopulation framework generally requires each population node in the network to have both a defined mosquito and human population size. Since the mosquito vectors we are interested in are anthropophilic, they tend to coexist with humans, so human population sizes and state distributions can be attributed to the same nodes at which mosquito populations are defined; however MGDrivE 2 also includes the possibility of human-only and mosquito-only nodes. Mosquito-only nodes could represent sites with only non-human animals from which mosquitoes bloodfeed, while human-only nodes could represent locations unsuitable for mosquitoes. As mosquitoes are able to move between nodes in the metapopulation, so can humans. This is an important factor to include, as human movement has been shown to drive the spatial transmission of mosquito-borne diseases such as dengue virus (Stoddard *et al.*, 2013).

### 2.3. Other extensions to inheritance, life history and landscape modules

Additional functionality has been included in the inheritance and life history modules of the MGDrivE framework since publication of version 1.0. The inheritance module is unchanged, and inheritance “cubes,” describing the distribution of offspring genotypes given maternal and paternal genotypes for a given genetic element, are usable in both versions. Several new inheritance cubes have been made available, including: a) homing-based remediation systems, including ERACR (Element for Reversing the Autocatalytic Chain Reaction) and e-CHACR (Eracing Construct Hitchhiking on the Autocatalytic Chain Reaction) (Gantz and Bier, 2016; Xu *et al.*, 2020), and b) newly proposed drive systems capable of regional population replacement, including CleaveR (Cleave and Rescue) (Oberhofer, Ivy and Hay, 2019) and TARE (Toxin-Antidote Recessive Embryo) drive (Champer *et al.*, 2020).

In the life history module, there are now two density-dependent functional forms to regulate population size - logistic and Lotka-Volterra - with the potential to add more. For mosquito vectors such as *Ae. aegypti* and *An. gambiae*, density-dependence is thought to act at the larval stage due to increased resource competition at higher larval densities (White *et al.*, 2011; Muriu *et al.*, 2013). The adult population size, *N*, is used to determine the carrying capacity of that habitat patch for larvae, *K*, which determines the degree of additional density-dependent mortality experienced by larvae at that patch. For the logistic model, the per-capita larval mortality rate is given by *μL* (1 + *L*(*t*)/*K*), where *μL* is the density-independent larval mortality rate, and *L*(*t*) is the total larval population size for the patch at time *t*. For the Lotka-Volterra model, the per-capita larval mortality rate is given by *μL* + *α L*(*t*), where *α* is the density-dependent term. Further details on these two density-dependent functions are provided in the S1 Text.

In the landscape module, movement through the network of population nodes is again determined by a dispersal kernel; however, due to the continuous-time nature of MGDrivE 2, movement between patches is described by a rate rather than a probability. The mathematical mapping between the rate matrix of MGDrivE 2 and the transition probability matrix of MGDrivE 1 is provided in the S1 Text.

### 2.4. Stochastic Petri net formulation

The most fundamental change from MGDrivE 1 to 2 is restructuring the model as a SPN (Haas, 2006). Adopting a SPN framework has several benefits. First, SPNs allow the mathematical specification of a model to be decoupled from its algorithmic implementation, allowing users to leverage extensive sampling algorithms from the physical and chemical simulation communities for efficient computation (Goss and Peccoud, 1998; Gillespie, 2007). Second, SPNs have a well-established and consistent formalism, allowing them to be readily understood and modified by anyone familiar with this (Gronewold and Sonnenschein, 1998). And third, SPNs are isomorphic to continuous-time Markov chains (CTMCs), meaning that model parameters can be time-varying, including Erlang-distributed aquatic stage durations and the pathogen EIP.

A Petri net is a bipartite graph consisting of a set of places, *P*, and a set of transitions, *T*. Directed edges or “arcs” lead from places to transitions (input arcs) and from transitions to places (output arcs). The set of arcs that connect places to transitions and transitions to places can be denoted by two matrices whose entries are non-negative integers describing the weight of each arc. The places define the allowable state space of the model; however, in order to describe any particular state of the model, the Petri net must be given a marking, *M*, which is defined by associating each place with a non-negative integer number of tokens. In the language of CTMCs, a marking, *M*, is referred to as a “state.” When a transition occurs, it induces a state change by “consuming” tokens in *M* given by the set of input arcs, and “producing” tokens in *M* according to the set of output arcs (Wilkinson, 2018). Each transition has a “clock process,” parameterized by a “hazard function” which defines that event’s current rate of occurrence. In MGDrivE 2, tokens represent an integer number of mosquitoes or humans, and the distribution of tokens (mosquitoes or humans) across states at time *t* defines a marking, *M*(*t*). A graphical representation of a Petri net for the mosquito life history module of MGDrivE 2 is depicted in Figure 3A, with a full description of the mathematical formalism provided in the S1 Text.

**Figure 3.**
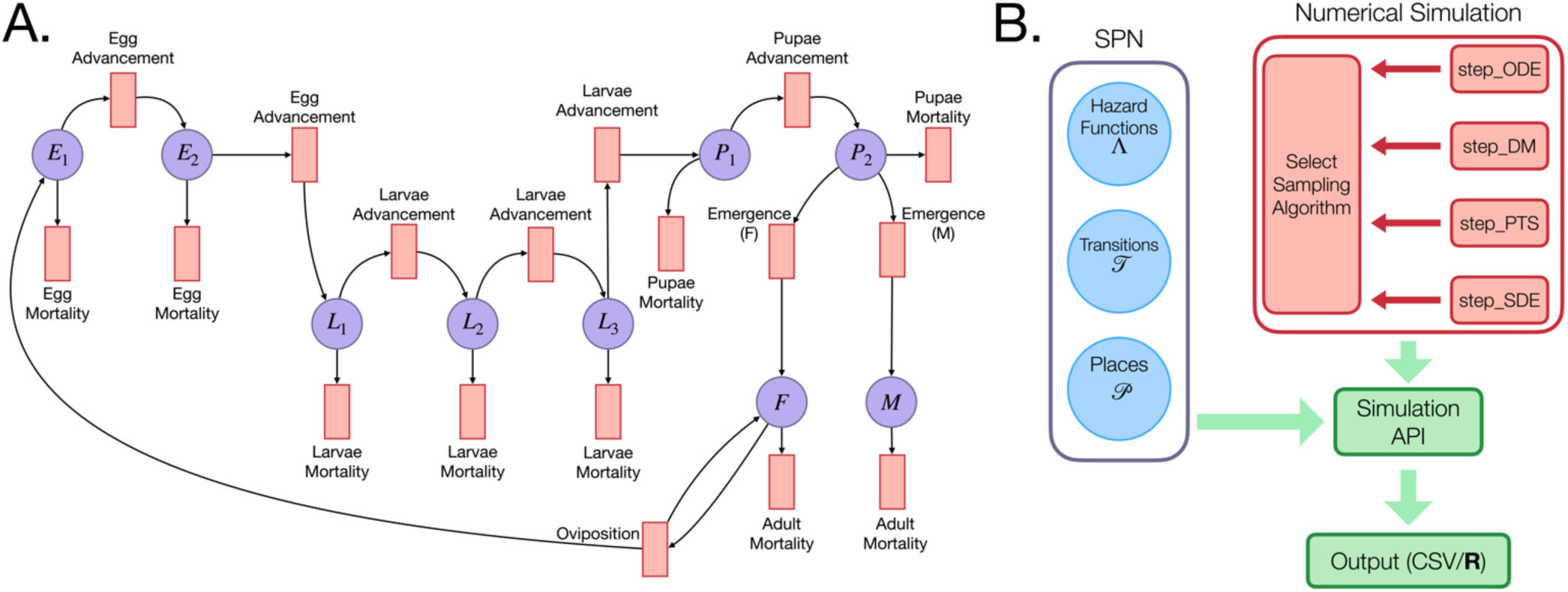
Stochastic Petri net (SPN) implementation of MGDrivE 2. **(A)** Petri net representation of the life history module. The set of purple circles corresponds to places, *P*, and red rectangles to transitions, *T*. This Petri net shows a model in which development times for the egg stage are Erlang-distributed with shape parameter *n* = 2, and for the larval stage are Erlang-distributed with shape parameter *n* = 3. Population dynamics are derived directly from this graph. E.g. The transition corresponding to oviposition has one edge beginning at F, meaning at least one female mosquito must be present for oviposition to occur. When oviposition occurs, a token is added to E1 (new eggs are laid) and a token is returned to F. **(B)** Conceptual representation of the SPN software architecture showing the separation between the model representation (blue circles) and set of sampling algorithms (red rectangles). These two components of the codebase meet at the simulation API, enabling users to match models and simulation algorithms interchangeably. Output may be returned as an array in R for exploratory work, or written to CSV files for large simulations.

The code that generates the Petri net is independent of the code that simulates trajectories from it. Once the Petri net is stored as a set of sparse matrices, it is passed to a simulation application program interface (API) which allows trajectories to be simulated as ordinary differential equations (ODEs), stochastic differential equations (SDEs), or CTMCs (Figure 3B). Each of these are referred to as “step” functions, but are not limited to discrete time steps; these functions are responsible for updating the model between time points where the user requests output to be recorded. The ODE step function provides a deterministic approximation and interfaces with the numerical routines provided in the “deSolve” R package (Soetaert, Cash and Mazzia, 2012). Three stochastic numerical routines are provided that treat the model as a continuous-time Markov process and provide different levels of approximation. The most straightforward method to sample trajectories is Gillespie’s direct method, which samples each event individually (Gillespie, 1977). While statistically exact, this is prohibitively slow for medium-to-large population sizes. Two approximate stochastic methods are provided that have been widely used in the chemical physics literature: i) a second order continuous SDE approximation known as the chemical Langevin equation (Gillespie, 1996), and ii) a fixed-step tau-leaping method (Gillespie, 2001). Both methods achieve substantial gains in computational speed at the expense of statistical accuracy. While the SDE approximation is often faster, tau-leaping retains the discrete character of the process it approximates and is usually the preferred technique. A full description of each of the numerical routines is provided in the S1 Text. In addition, we demonstrate how a user can write a custom simulation algorithm and incorporate it within the MGDrivE 2 codebase in the “Advanced Topics” vignette available at https://marshalllab.github.io/MGDrivE/docs_v2/articles/advanced_topics.html.

## 3. Results

To demonstrate how the MGDrivE 2 framework can be used to initialize and run a simulation of the spread of a gene drive system through a metapopulation with time-varying model parameters, including its implications for vector-borne pathogen transmission, we have provided vignettes with the package, available via installation from our repository at https://github.com/MarshallLab/MGDrivE/tree/master/MGDrivE2 and additional examples and information on Github at https://marshalllab.github.io/MGDrivE/docs_v2/index.html. The vignettes provide extensive examples of how to use the software, including advanced features such as implementing custom time-varying rates and numerical simulation algorithms. They consist of a set of five “core” manuals that describe how to simulate population genetics and dynamics for a mosquito-only population and metapopulation, then how to incorporate SEI-SIS Ross-Macdonald malaria transmission dynamics in a population with humans included, and finally how to incorporate SEI-SEIR arbovirus transmission dynamics. Following these are three “advanced” manuals that introduce: i) how to process and analyze output from simulations that write to CSV files, ii) how users can write custom time-varying hazard functions, and iii) how a user might implement their own numerical simulation routine, using an explicit Euler method for ODEs as an example.

Here, we describe the application of the package to model the release of a population replacement gene drive system designed to drive a malaria-refractory gene into an *An. gambiae* mosquito population with seasonal population dynamics and transmission intensity calibrated to a setting resembling the island of Grand Comore, Union of the Comoros. The gene drive system resembles one engineered in *An. stephensi* that is integrated into the *kynurenine hydroxylase* gene and includes a recoded copy of that gene that rescues its function (Adolfi *et al.*, no date). This design selects against resistance alleles that interrupt its function. Four alleles are considered: an intact homing allele (denoted by “H”), a wild-type allele (denoted by “W”), a functional, cost-free resistant allele (denoted by “R”), and a non-functional or otherwise costly resistant allele (denoted by “B”). Full details of the inheritance dynamics are provided in Adolfi *et al.* (2020) and model parameters are summarized in Table S1.

The life history module is parameterized with typical bionomic parameter values for *An. gambiae* (Table S1), including mean-variance relationships describing the development times of juvenile life stages (Bayoh and Lindsay, 2003). The carrying capacity of the environment for larvae is a function of recent rainfall, and the adult mortality rate is a function of temperature. Remotely sensed rainfall data for Grand Comore was obtained from the ERA5 dataset (https://www.ecmwf.int/en/forecasts/datasets/reanalysis-datasets/era5), and a mathematical relationship adapted from White *et al.* (White *et al.*, 2011) was used to translate this to larval carrying capacity, assuming that half of the island’s carrying capacity was provided by permanent breeding sites (e.g. large cisterns) and half was provided by recent rainfall. Temperature data for Grand Comore was also obtained from the ERA5 dataset, and adult mortality was derived using methods described by Mordecai *et al.* (Mordecai *et al.*, 2019). Both climatological time series covered the ten year period beginning January 1, 2010. For the purpose of this demonstration, Grand Comore was treated as a single randomly mixing population, although simulations involving a more detailed landscape module are included in the vignettes.

The epidemiology module is parameterized with typical parameter values for *Plasmodium falciparum* transmission (Table S1), human population size and life expectancy parameters from the National Institute of Statistics and Demographic Studies, Comoros (INSEED, 2015), and is calibrated to local malaria prevalence estimates from the Malaria Atlas Project (Pfeffer *et al.*, 2018). This calibration was achieved by multiplying the carrying capacity time series by a constant such that the average adult female mosquito population over a year sustained malaria transmission in the human population at the estimated local prevalence. Finally, we caution that these simulations are merely intended to demonstrate the software’s capabilities and that, while the simulations are parameterized with data from Grand Comore, they are not intended to provide an accurate forecast of local gene drive mosquito dynamics, or to imply approval of the intervention by the local population and regulatory agencies.

### 3.1 Simulation workflow

The code for this simulation is available at https://github.com/MarshallLab/MGDrivE/tree/master/Examples/SoftwarePaper2. We begin by loading the MGDrivE 2 package in R, as well as the package for the original MGDrivE simulation, which provides the inheritance cubes required for simulation of genetically-stratified mosquito populations. Next, we define model parameters, including the bionomic parameters of *An. gambiae* s.l., and demographic and epidemiological parameters specific to Grande Comore. To parameterize time-varying adult mosquito mortality (hourly) and larval carrying capacity (daily), we load CSV files containing those data as time series for the ten year simulation period. We then use the base “stepFun()” function in R to create an interpolating function of those time-series data that will return a value for any time within the simulation period, which is required for calculation of hazard functions. More sophisticated interpolating functions, such as splines, may also be used. We also specify the inheritance cube at this point, as the number of modeled genotypes and distribution of offspring genotypes for given parental genotypes will be used to build the Petri net.

Next, we use functions from MGDrivE 2 to create the “places” and “transitions” of the Petri net, which are stored as lists in R and then converted into a sparse matrix representation used in the simulation code. Epidemiological dynamics and states are coded automatically by calling the functions that create the Petri net. In this case, “spn_P_epiSIS_node()” and “spn_T_epiSIS_node()” will generate the places and transitions for a single node model with SEI-SIS mosquito and human malaria transmission dynamics. Each transition has a tag that specifies the hazard function it requires. Following that, we write custom time-varying hazard functions for adult mosquito mortality and larval mortality (a function of carrying capacity). We provide a guided walkthrough of how a new user might write their own time-varying hazard function in the vignette “Simulation of Time-inhomogeneous Stochastic Processes.” Once the vector of hazard functions has been stored (as a list), we create the data frame that stores the times, genotypes, sex, and size of each release event.

With the construction of all model components necessary for the simulation, we call the simulation API which handles the details of simulating trajectories from the model. In this case, we chose the tau-leaping algorithm to sample stochastic trajectories, and to record output on a daily basis. MGDrivE 2 allows users to choose how model output is reported back - for exploratory or smaller simulations, users may return output directly to R as an array; however for larger simulations, it is often preferable to write directly to CSV files due to memory considerations, and MGDrivE 2 has sophisticated functions to both specify CSV output and process completed simulations.

### 3.2. Entomological population dynamics

In Figure 4, we display a potential visualization scheme produced in Python for the simulations described above. The code to produce this visualization is available at https://github.com/Chipdelmal/MoNeT/tree/master/DataAnalysis/v2 (note that MGDrivE 2 code does not depend on Python). Figure 4A displays the climatological time-series data - temperature in magenta and rainfall in blue - which were used to calculate time-varying adult mosquito mortality rate and larval carrying capacity, respectively. The total adult female population size averaged over 100 stochastic runs is shown in green. This is relatively consistent throughout the year due to moderate seasonal changes in temperature in the tropical climate of the Comoros and the presence of permanent breeding sites such as cisterns throughout the island. Figure 4B displays allele frequencies for adult female mosquitoes over the simulation period. After eight consecutive weekly releases of 10,000 male mosquitoes homozygous for the drive allele (HH) three years into the simulation, we see the drive allele (H) spread to high frequency in the population, the wild-type allele (W) be completely lost, and the in‐frame resistant allele (R) accumulate to a small but noticeable extent. This occurs due to the drive of the H allele, and because the R allele is generated at a low rate and has neither a fitness cost nor benefit relative to the H and W alleles. The out-of-frame or otherwise costly resistant allele (B) initially rises in frequency more quickly than the R allele due to its higher generation rate, but declines in frequency once there are no more W alleles to cleave due to its inherent selective disadvantage.

**Figure 4.**
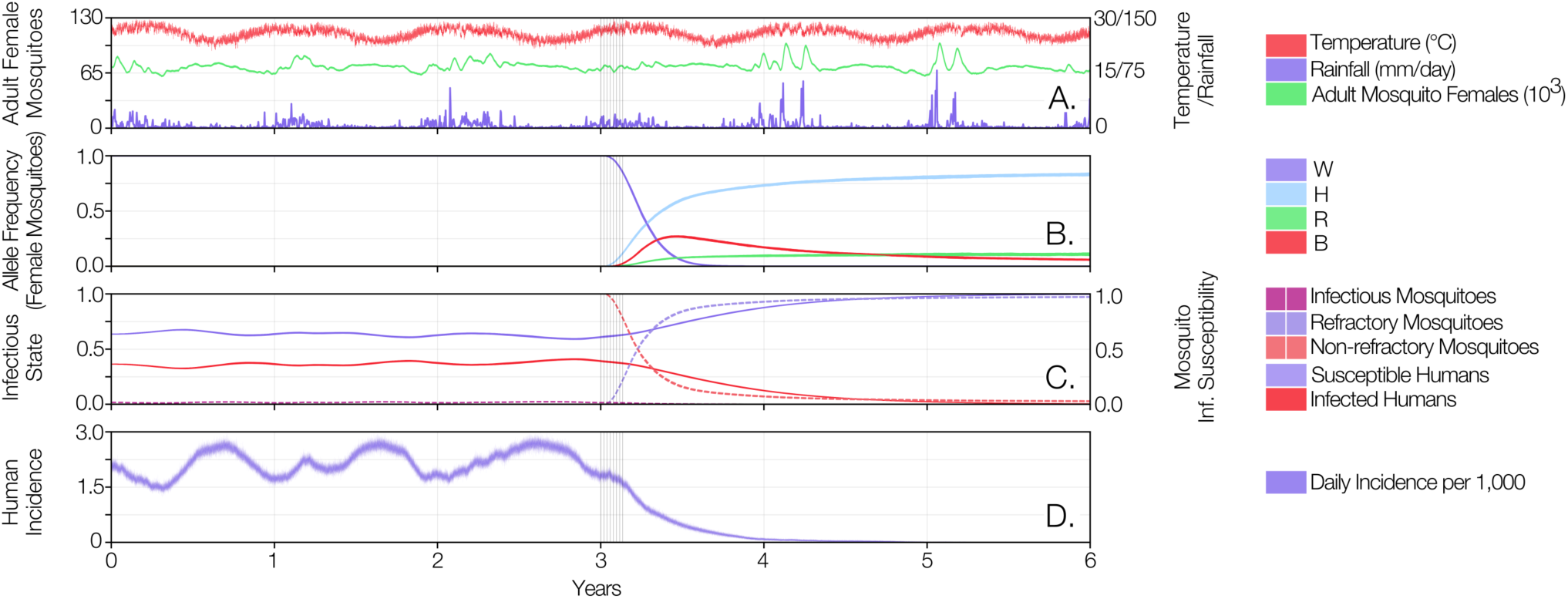
Example MGDrivE 2 simulations for a population replacement gene drive system designed to drive a malaria-refractory gene into an *An. gambiae* s.l. mosquito population with seasonal population dynamics and transmission intensity calibrated to a setting resembling the island of Grand Comore, Union of the Comoros. The gene drive system resembles one recently engineered in *An. stephensi* (Adolfi *et al.*, 2020) for which four alleles are considered: an intact homing allele and malaria-refractory gene (denoted by “H”), a wild-type allele (denoted by “W”), a functional, cost-free resistant allele (denoted by “R”), and a non-functional or otherwise costly resistant allele (denoted by “B”). Model parameters describing the construct, mosquito bionomics and malaria transmission are summarized in Table S1. **(A)** Climatological time-series data - temperature in red and rainfall in purple - that were used to calculate time-varying adult mosquito mortality rate and larval carrying capacity, respectively. The resulting adult female population size is shown in green. **(B)** Allele frequencies for adult female mosquitoes over the simulation period. Grey vertical bars beginning at year three denote eight consecutive weekly releases of 10,000 male mosquitoes homozygous for the drive allele (HH). **(C)** Spread of the malaria-refractory trait through the female mosquito population, and consequences for mosquito and human infection status. Following the release of the drive system at year three, the proportion of refractory female mosquitoes (dotted light purple line) increases and the proportion of infectious mosquitoes (dotted dark purple line) declines. As humans recover from infection and less develop new infections, the *P. falciparum* parasite rate (solid red line) declines until it reaches near undetectable levels by year five. **(D)** Human malaria incidence is halted by the beginning of year four.

### 3.3. Epidemiological dynamics

The gene drive system we consider includes a malaria-refractory gene that results in complete inability of mosquitoes to become infected with the malaria parasite, whether present in either one or two allele copies. In Figure 4C, we depict the spread of the malaria-refractory trait through the female mosquito population, and the consequences this has for mosquito and human infection status. Prior to the release, we see that infection prevalence in humans (*P. falciparum* parasite rate, PfPR) is mildly seasonal, with the proportion of infected humans (solid red line) waxing and waning in response to the fluctuating mosquito population size (green line in Figure 4A). The proportion of infectious female mosquitoes (dotted dark purple line) oscillates in synchrony with the proportion of infected humans; but at a much lower proportion due to the short mosquito lifespan and the fact that most mosquitoes die before the parasite completes its EIP. Following the release of the drive system and refractory gene at year three, the proportion of refractory female mosquitoes (dotted light purple line) increases and, consequently, the proportion of infectious mosquitoes declines. As humans recover from infection and less develop new infections, the PfPR declines until it reaches near undetectable levels by year five. Lastly, Figure 4D depicts human malaria incidence, measured as the number of new infections per 1,000 humans per day. Stochastic variation in this model output is more pronounced due to the small number of incident cases relative to the total population. Incidence is halted by the beginning of year four, but PfPR takes almost a year longer to approach zero as infected humans clear parasites.

## 4. Future directions

We are continuing development of the MGDrivE 2 software package and welcome suggestions and requests from the research community regarding future directions. The field of gene drive research is moving quickly, and we intend the MGDrivE 2 framework to serve as a flexible tool to address exploratory, logistical and operational questions regarding genetics-based control systems for mosquito disease vectors. This includes exploratory modeling of novel genetic constructs, assessment of candidate constructs against TPPs and PPCs, and field trial planning as constructs progress through the development pipeline. Future functionality that we are planning includes: i) modeling of mosquito traps to address questions related to monitoring and surveillance, and ii) more detailed epidemiological models addressing phenomena important to malaria and arbovirus transmission - for instance, dengue models that incorporate multiple serotypes with temporary cross-protective immunity and complications related to antibody-dependent enhancement (Wearing and Rohani, 2006), and malaria models that incorporate age-structure, immunity, asymptomatic infection and superinfection (Griffin *et al.*, 2010).

Additionally, we are exploring numerical sampling algorithms that can increase computational efficiency and speed, facilitated by separation of model specification and simulation in the software. The complexity of models that can be developed in MGDrivE 2 means that sensitivity analyses can become extremely computationally intensive, and the ability of the SPN framework to leverage efficient algorithms in these circumstances will be highly valuable. We also continue to be interested in developing a corresponding individual-based model capable of efficient modeling when the number of possible states exceeds the number of individuals in the population - for instance, for multi-locus systems such as daisy-drive (Noble *et al.*, 2019) and multiplexing schemes in which a single gene is targeted at multiple locations to reduce the rate of resistance allele formation (Prowse *et al.*, 2017), and for epidemiological models in which age structure, immunity and mosquito biting heterogeneity become prohibitive for population models (Griffin *et al.*, 2010).

## Acknowledgements

This work was supported by a DARPA Safe Genes Program Grant (HR0011-17-2-0047) and funds from the UC Irvine Malaria Initiative and Innovative Genomics Institute awarded to J.M.M.

## Author contributions

S.L.W. and J.M.M. conceived the project. S.L.W. led MGDrivE 2 development and J.B.B. and H.M.S.C. contributed substantially to core development. A.J.D. contributed to development of the SPN framework. T.M.L. contributed to the ecological component. S.L.W. and J.M.M. wrote the first draft of the manuscript. S.L.W. and J.B.B. wrote the vignettes. All authors revised the manuscript and approved for publication.

## Data availability statement

MGDrivE 2 is available at https://github.com/MarshallLab/MGDrivE/tree/master/MGDrivE2. The source code is under the GPL3 License and is free for other groups to modify and extend as needed. Mathematical details of the model formulation are available in the S1 Text, and documentation for all MGDrivE 2 functions, including vignettes, are available at the project’s website at https://marshalllab.github.io/MGDrivE/docs_v2/index.html. To run the software, we recommend using R version 2.10 or higher.

## 1. Lifecycle Model

The lifecycle model is similar to the discrete time ecology module used in **MGDrivE** (Sánchez C. et al. 2019). Major differences include the switch to continuous time and replacement of fixed, constant delays with Erlang distributed delays in aquatic life stages. This change means that, whereas **MGDrivE**’s deterministic model was formulated as a set of delay difference equations, **MGDrivE 2**’s deterministic model is a set of ordinary differential equations (ODEs) (using the “linear chain trick” to simulate Erlang-distributed delays, (Hurtado and Kirosingh 2019)).

Similar to **MGDrivE**, the lifecycle model includes egg (E), larval (L), and pupal (P) aquatic stages. Upon emergence from P, adult mosquitoes are assigned a sex, the probability of which may depend on genotype. Upon emergence, females (F) become mated in the presence of male mosquitoes (M), and oviposit at an age-independent (though possibly time-dependent) rate until they die. If there are no adult males, newly emerging females transfer to an unmated adult female (U) compartment, where they remain until death or successful mating if males become available.

The system of ODEs describing the deterministic lifecycle model are solved at their non-trivial equilbrium to provide initial conditions for simulations in **MGDrivE 2**. These ODEs are a limiting case of the stochastic continuous-time Markov chain (CTMC) model, when populations are large (for technical conditions, see (Kurtz 1970)). In our presentation of the ODEs and their equilibrium solutions, we ignore indexing by genotype because, for most simulations, the equilibrium solution corresponds to a baseline scenario prior to releases of modified mosquitoes, where mosquito populations are composed solely of wild-types. These equations also ignore indexing by node. We use (*n*_*E*_, *n*_*L*_, *n*_*P*_) to denote the number of sub-stages in each aquatic stage (the Erlang shape parameter). Subscript *i* refers to any particular sub-stage, such that eggs are denoted *E*_*i*_, larvae as *L*_*i*_, pupae as *P*_*i*_, and *N*_*F*_, *N*_*M*_ the mated adult female and male populations, respectively. Because the non-trivial equilibrium will have zero unmated females, there is no *N*_*U*_ compartment.

Shape and rate parameters for the Erlang-distributed delays can be constructed as follows. Consider a random delay with mean 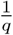 and variance 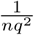, where *n* is an integer. Such a random delay can be assumed to follow an Erlang distribution, and one way to construct a model of this system is to build a linear system of *n* bins, where the rate of transfer from the *i* − 1*^th^* to *i*^*th*^ bin is given by *qn*. In a deterministic model, this will be a linear system of ODEs and, for a stochastic model, a CTMC.

We present the life history model with two different parameterizations of larval density dependence, which we call the “Lotka-Volterra” and “Logistic” versions, in reference to ecological theory. Both sets of equations are available in the code, and are provided as an example of how to use different functional forms of rate equations with the same Petri net structure.

### 1.0.1. Lotka-Volterra Density-Dependent Equations

This set of equations uses a linear form of per-capita density-dependent mortality for the larval instar stages that corresponds to the functional form assumed by (Hancock and Godfray 2007).

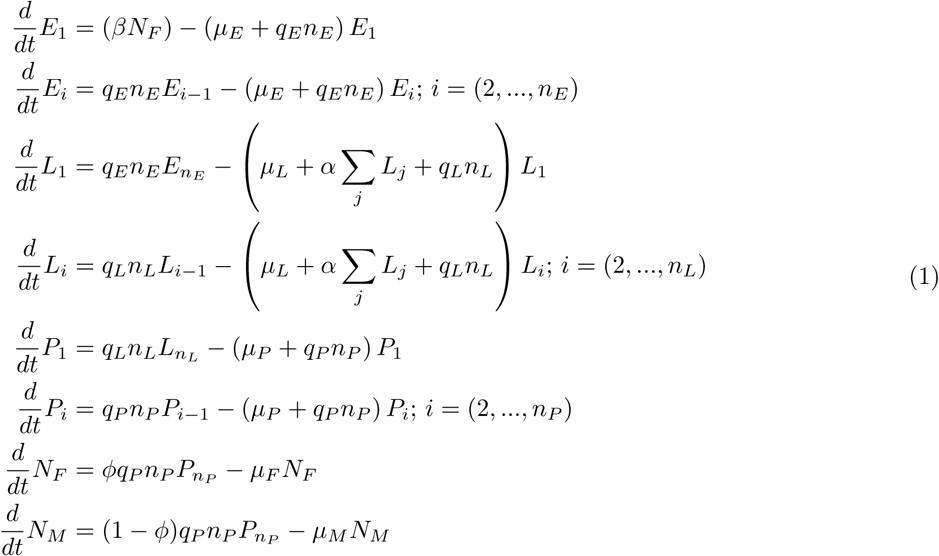

In this set of equations, the parameter *α* represents increased mortality rates that occur as a function of crowding, and has units of time^*−*1^area^2^.

To solve the model with linear density-dependence at equilibrium, we assume that the equilibrium number of adult female mosquitoes, 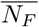 is known, and that all rate constants, with the exception of *α* are also fixed; we then solve for all remaining state variables plus *α*, giving a system of 2 + *n*_*E*_ + *n*_*L*_ + *n*_*P*_ equations and the same number of unknowns.

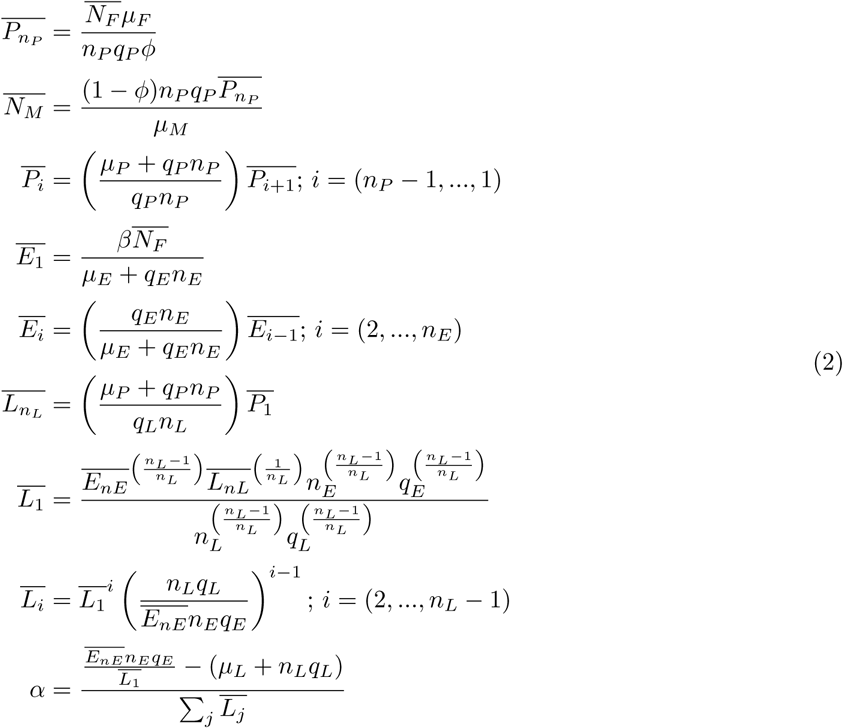

### 1.0.2. Logistic Density-Dependent Equations

This set of equations uses a rational form of per-capita density-dependent mortality for the larval instar stages that uses a carrying capacity *K* parameterization (equivalent to the logistic model in ecology).

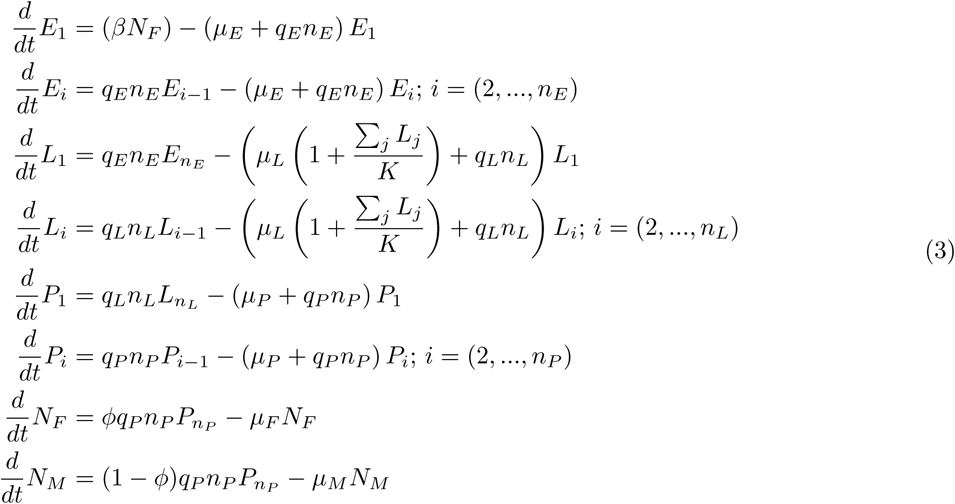

In this parameterization of density-dependent mortality, *µ*_*L*_ is the natural mortality rate of larvae without any effects of resource depletion or competition (because when Σ_*j*_ *L*_*j*_ is small, the mortality is approximately *µ*_*L*_).

To solve the model at equilibrium, we assume that the equilibrium number of adult female mosquitoes, 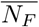 is known, and that all rate constants, with the exception of *K*, are also fixed; we then solve for all remaining state variables plus *K*, giving a system of 2 + *n*_*E*_ + *n*_*L*_ + *n*_*P*_ equations and the same number of unknowns.

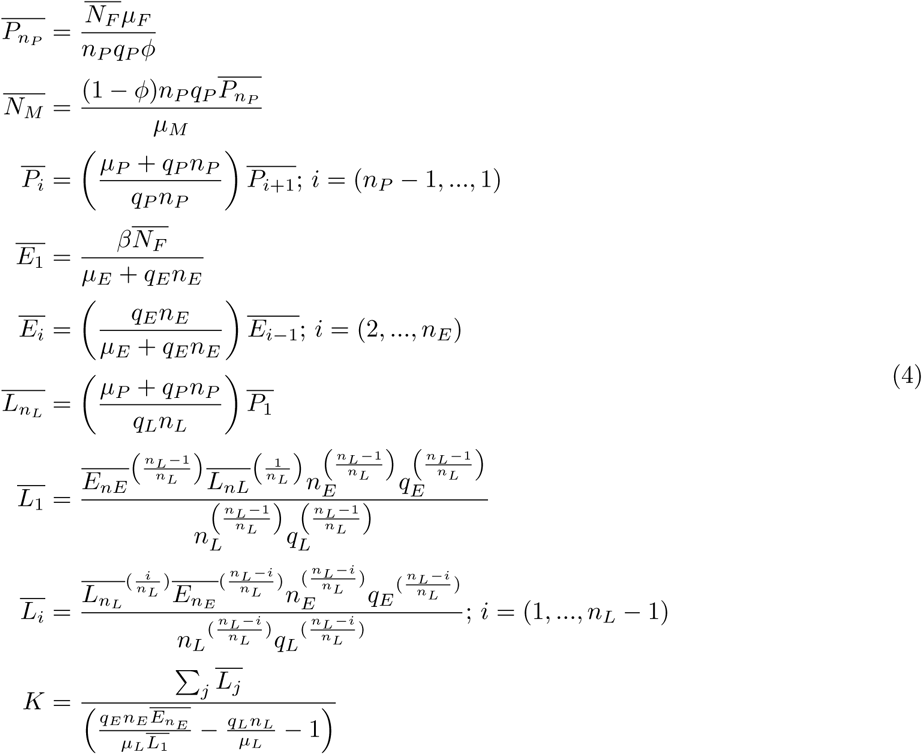

In fact, because both of these per-capita density dependent rates of mortality are linear functions in the number of larvae present (such that the overall mortality is quadratic in the number of larvae), at equilibrium the parameters *α* and *K* are related by the simple expression:

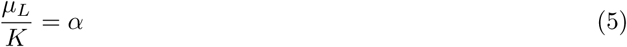

### 1.1. Parameters

Due to the continuous-time model structure as well as reformulating the fixed delays of **MGDrivE** as Erlang-distributed random delays, parameters used in **MGDrivE** cannot be directly “plugged-in” to **MGDrivE 2** simulations. In this section we discuss how to parameterize **MGDrivE 2**, and discuss similarities and differences with those in (Marshall, Buchman, Akbari, et al. 2017; Sánchez C. et al. 2019). We note that there will be certain mathematical artifacts which prevent a one-to-one mapping between the two models due to the change between a lagged discrete-time Markov chain (DTMC) to continuous-time Markov chain (CTMC) model formulation. For more details on how these arise and their effects, please consult (Fennell, Melnik, and Gleeson 2016; Allen 1994).

#### 1.1.1 Aquatic Survival

Let the probability to survive any aquatic state *x* ∈ {*E*, *L*, *P*} be *θ*_*x*_. In **MGDrivE**, these were given as:

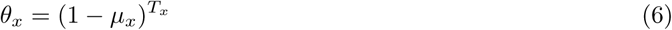

In **MGDrivE 2**, the aquatic state is broken in *n*_*x*_ substages to produce an overall Erlang-distributed dwell time, *τ*. The Erlang distribution has shape parameter *n*_*x*_ and rate parameter *n*_*x*_*q*_*x*_, where 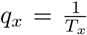 therefore 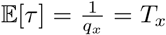 and 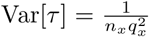. Because the dwell time *τ* is a random variable the probability of survival is expressed:

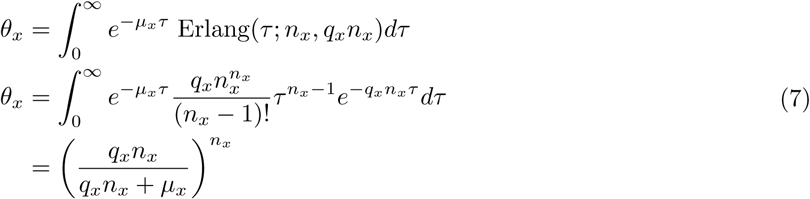

If we wanted to match survival probabilities between the two models, we just consider *µ*_*x*_ in equation 7 to be an unknown and solve for it:

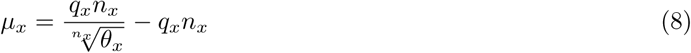

Note that we can arrive at the solution from equation 7 by considering not a single random variable *τ* but rather the random variables *X, Y*, where the latter is the time to death, if death were to occur, and the former is time to advancement out of stage *x*, were advancement to occur. Then we want the probability that *X < Y*:

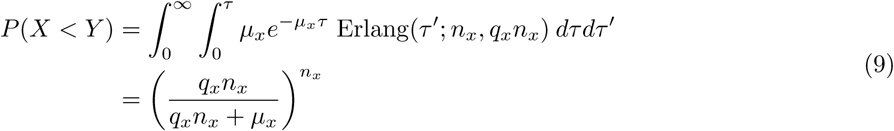

Intuition behind the solution may be acquired if we take literally the interpretation of the Erlang distribution as being used in the “linear chain trick”; in this case at each substage the overall probability of survival is 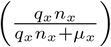. Because there are *n*_*x*_ substages, the total survival probability is the product of the *n*_*x*_ stages.

#### 1.1.2. Population Growth Rate

In **MGDrivE**, the intrinsic population growth rate *R*_*m*_ was defined as “equal to the rate of female egg production multiplied by the life expectancy of an adult mosquito multiplied by the proportion of eggs that will survive through all of the juvenile life stages in the absence of density-dependence” (Marshall, Buchman, Akbari, et al. 2017).

In **MGDrivE** it had units of mosquito^−1^day^−1^:

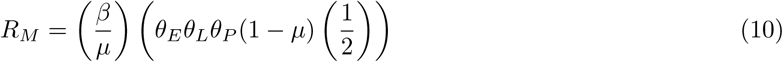

It is essentially the same in **MGDrivE 2**, using the form of *θ*_*x*_ from equation 6:

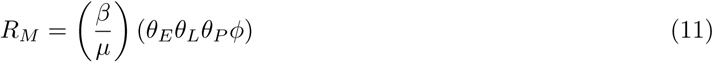

Note however, the absence of the 1 − *µ* term; this is because equation 11 is a continuous time rate; adults are available to oviposit immediately upon emergence, so there is no need extra mortality between emergence and adulthood (Deredec, Godfray, and Burt 2011).

#### 1.1.3. Parameterization from Growth Rates

In **MGDrivE** the model was typically parameterized such that equilibrium solutions were available in closed form. The assumptions, outlined in the supplemental information of (Marshall, Buchman, Akbari, et al. 2017) and based on the model of (Deredec, Godfray, and Burt 2011), are that *µ*_*E*_ = *µ*_*L*_ = *µ*_*P*_; that is, the density-independent mortality of each aquatic stage is the same. In the absence of density-dependent effects, the total probability of surviving the aquatic stages was 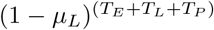. Combined with knowledge of the generation time *g*, the daily population (geometric) growth rate *r*_*M*_, and the per-generation geometric growth *R*_*M*_ = (*r*_*M*_)*^g^*, *µ*_*L*_ could be found in closed form, and from that the remaining unknowns, *α* and *L*_*eq*_ (strength of density dependence and equilibrium larval population) could also be solved in closed form. The relevant equations were S51-S55 in (Sánchez C. et al. 2019).

In **MGDrivE 2** we want to be able to solve for equilibria under similar assumptions of equal density-independent mortality across aquatic stages. Let us define *µ*_*A*_ = *µ*_*E*_ = *µ*_*L*_ = *µ*_*P*_ so we seek a solution to the unknown constant aquatic stage mortality *µ*_*A*_. We first expand Equation 11 in terms of Equation 7:

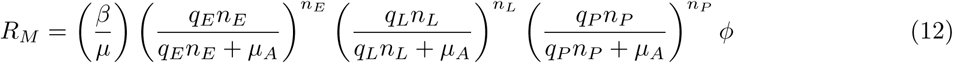

Here, *R*_*M*_ is the per-generation growth rate; in **MGDrivE** it was (*r*_*M*_)^g^. However, because **MGDrivE 2** is a continuous time model, the equations for geometric growth are not appropriate. The equivalent infinitesimal rate of growth is log (*r*_*M*_) such that 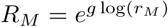, which is the left hand side of Equation 12.

Finding closed form solutions to this equation is difficult because of the additional terms introduced by the Erlang delays. Therefore we use Newton’s method in **R** (with `uniroot`) to numerically solve for *µ*_*A*_. The input to the function is the daily growth rate *r*_*M*_ and biological parameters *β,μ,q*_*E*_,*n*_*E*_*q*_*L*_,*q*_*P*_,*n*_*P*_,*ϕ*, and it returns the value of *µ*_*A*_ such that the following equation holds:

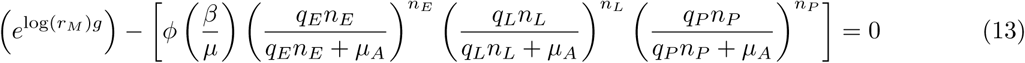

#### 1.1.4 Lifespan Modification

In **MGDrivE**, the genotype-specific parameter *ω* was used to reduce lifespans of non wild-type organisms due to fitness costs associated with the homing cassette, or intentional fitness reduction. However, for **MGDrivE** 2 we generalize this to modify lifespans in either direction, relative to wildtype, because experimental data showed evidence of in some cases substantial lifespan *increases* from driving certain genetic material in model organisms (Kandul et al. 2019).

Because wildtype lifespans (*x*) were geometrically distributed random variables, and daily survival was given by (1 − *µ*)*ω*, one could solve for a reduced lifespan, *y* < *x*, by noting the daily mortality probability can be written *p*= 1 − ((1 − *μ*)*ω*) = 1 − *ω* + *μω*. Then note that the mean lifespan is 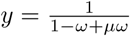. Thus to solve for *ω*, we solve for the root of the equation 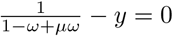, where only *ω* is unknown.

In **MGDrivE 2** adult lifespans are exponentially distributed random variables. To change the lifespan *y* (no longer restricted to *y* < *x*, *y* may be any positive number), consider modifying the mortality hazard by the factor *ω*. To find *ω* we just solve the following equation:

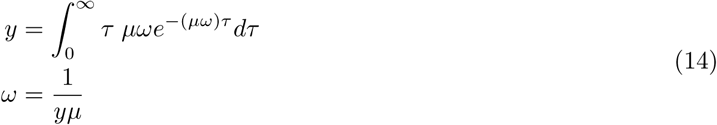

#### 1.1.5 Movement

In the continuous time model, mosquito movement is given by a rate of movement from each node *i* to all other nodes *j* ≠ *i*. When parameterizing these rates, we need to take into account that they will be a function of the total probability of a mosquito to leave its natal habitat *i* over its lifetime, *P*. Given that adult mosquitoes are subject to mortality with rate *µ*, one can solve for the rate of movement out of node *i* to anywhere (*δ*) as follows:

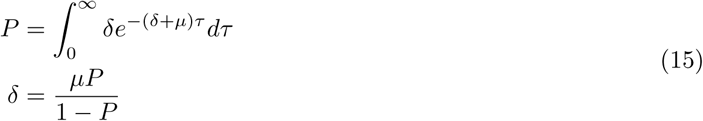

Then, if from node *i*, we have some vector of movement probabilities {*π*_*ij*_}_*j≠i*_ which give the probability to move from *i* to *j* conditional on leaving *i*, we can set the movement hazard between to be *δπ*_*ij*_ so we get the right lifetime leaving probability. The new vector of movement *hazards* is {*δπ*_*ij*_}_*j≠i*_

### 1.2 Genetic Inheritance & Modification

Because **MGDrivE 2** builds upon our previous work (Sánchez C. et al. 2019), the data structure used in **MGDrivE** to store all probabilities, fitness costs, etc. related to genotypes (the “inheritance cube”) is compatible with **MGDrivE 2**. In fact, we use the same cubes developed in that model and do not introduce any new cubes in this text.

While data structures can be reused, the fitness modifiers that describe the effect of inherited genotype on the life-history have a slightly different interpretation. In **MGDrivE**, genotype specific multipliers applied to daily probabilities, they had to be bounded in order to prevent nonsensical parameter values. If *P* represents a wild-type daily survival probability, for example, the modifier *ω* must be 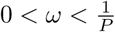. Because **MGDrivE 2** is parameterized directly in terms of hazards, the genotype-specific effects can be any positive real number. This has the added benefit of making them amenable to parameterization directly from survival analysis routinely preformed on biological lab experiments which often estimate relative hazards (Kandul et al. 2019).

## 2 Epidemiological Dynamics

To introduce epidemiological dynamics in **MGDrivE 2**, we use the SEI-SIS coupled model of mosquito and human infection dynamics as the basic model (Figure S1), as more complex vector-host models tend to be modifications of the basic form. This type of model is known as the Ross-Macdonald model in mathematical epidemiology (Martcheva 2015; Smith and McKenzie 2004). **MGDrivE 2** also supports SEI-SEIR models, which we introduce briefly later.

Here *S*_*H*_ refers to susceptible humans, *I*_*H*_ to infected/infectious humans, *S*_*V*_ to susceptible mosquitoes, (*E*_*V*,1_,…, *E*_*V,n*_) to incubating mosquitoes, and *IV* to infectious mosquitoes. Because only mated adult females undergo gonotrophic cycles which require bloodfeeding, infection dynamics are only present in *F*.

To investigate the dynamics of the model, it is important to focus on events (transitions) that change state, and the rate at which they occur. For epidemiological dynamics, the two primary events are mosquito to human transmission and human to mosquito transmission, each driven by a Poisson process. Computation of the rate with which each process occurs in time depends on the per-capita force of infection (FOI) terms: *λ*_*H*_ and *λ*_*V*_, the rates at which any particular susceptible human gets infected and moves to the infected class, and the rate at which any particular susceptible mosquito gets infected and moves to the incubating class, respectively. When multiplied by the total numbers of susceptible humans or susceptible mosquitoes, respectively, we arrive at the correct rate for the Poisson processes.

**Figure S1:**
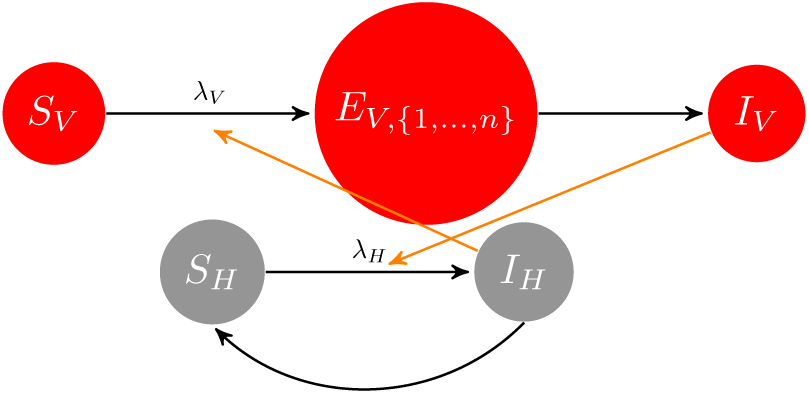
SEI-SIS pathogen transmission system; orange arrows denote the contribution of each species to the force of infection term on the other.

### 2.1. Transmission Terms

The function *λ*_*H*_ is the per-capita FOI on susceptible humans, such that *λ*_*H*_*S*_*H*_ is the total rate at which infection in the human population occurs in the deterministic model, or the intensity of the Poisson process for human infections in the stochastic model. Using Ross-Macdonald parameters as in (Smith and McKenzie 2004), 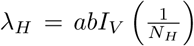. This is because, if *a* is the human biting rate, *b* mosquito-human transmission efficiency, then the total number of infectious bites produced by the mosquito population is *abIV*. Assuming uniform biting on humans, any particular person has probability 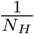 of being bitten, so 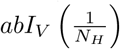 is the per-capita FOI. Multiplication by *S*_*H*_, the total number of humans, gives the total rate of infection in the human population.

The per-capita FOI in the mosquito population is *λ*_*V*_ *S*_*V*_. The FOI on susceptible mosquitoes is written as 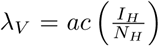. Again, *a* is the human biting rate, *c* is the human-mosquito transmission efficiency, and 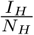 is the probability that a bite lands on an infectious human. Multiplying by the total susceptible vector population, *S*_*V*_, gives the total rate of infection in the mosquito population.

### 2.2. Deterministic Approximation

In this section we describe how to develop a mean-field approximation given by a system of ODEs to the stochastic SEI-SIS system.

#### 2.2.1. Mean-field Approximation of Human Stochastic Dynamics

Here we describe in detail the method used to approximate the stochastic CTMC model of infection dynamics in the human population, as its state space is smaller and the same methods can be used for the larger mosquito dynamics. The methods follow those presented in (Wilkinson 2006). Because we are only considering the human population, we drop the *H* superscript on state variables.

If we consider the mosquito population to be constant, as it would be at dynamic equilibrium for the deterministic model, then *λ*_*H*_ will be a constant and we can decouple the human SIS dynamics from the mosquito SEI system. We show a diagram of the human only dynamics in Figure S2.

**Figure S2:**
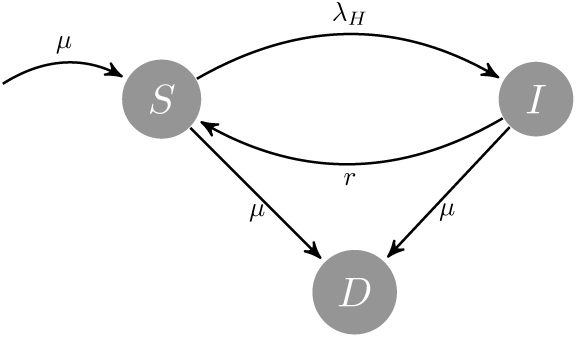
Susceptible-infected-susceptible (SIS) human infection dynamics

One way to analyze a CTMC is to derive the *Chapman-Kolmogorov equations* of the process. These equations give the conditional probabilities to transition to any ending state from a given starting state, over some time interval. For example if at time *t* ≤ *t*′ the process (represented by *X*(*t*)) is in a state *x*, then the probability to jump to any state *x*′ is **P**(*X*(*t + t′* = *x*′|*X*(*t*) = *x*), where **P** is a distribution over future states. It is often easier to work with the differential form of these equations, where the derivative is taken with respect to time, giving a system of differential equations that describes the time evolution of the Markov transition kernel over state space. Taking the derivative of these equations will involve expanding **P**(*t* + *δt*), which if expanded as **P**(*t*)**P**(*δt*), leads to the linear system of ODEs known as the Kolmogorov forward equations (KFE).

The CTMC is a process which jumps between points in state space (*S,I*) → (*S*′,*I*′). Put another way, the joint density must be understood as giving the probability to transition between all unique pairs of ways a number of people can be either susceptible or infectious, when birth, death, infection, and recovery can change state. Taking into account all events that can cause the system state at time *t* to change, we can derive the KFE as:

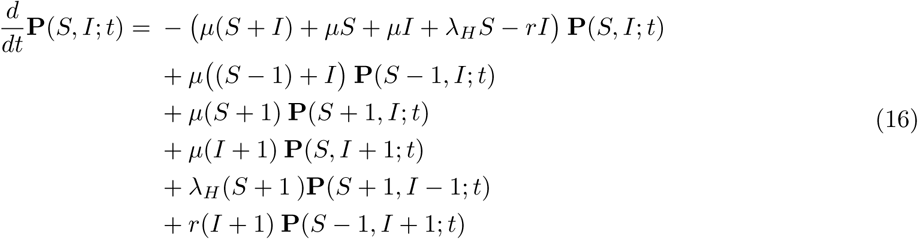

The KFE can be manipulated to derive the deterministic approximation to the CTMC. Because we assume constant *λ*_*H*_, the hazard/rate functions for each event are all first order in the state variables, the deterministic approximation will accurately describe the expected value of the stochastic model.

To go about this, we first construct the *stoichiometry matrix* **S**_*u×v*_, where *u* is dimension of the state space, and *v* is the number of unique events in the process. As the dimension of both state and event spaces are small, we can easily write this as:

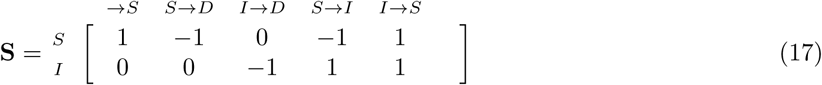

To begin developing our deterministic approximation, we take the time-derivative of the expectation of our state vector at time *t*, denoted as *X*(*t*) (full derivation in (Wilkinson 2006)). We also define *h*(*X*(*t*)) as a *v*-dimensional column vector of rates/intensities of each event at that point *x* in state space.

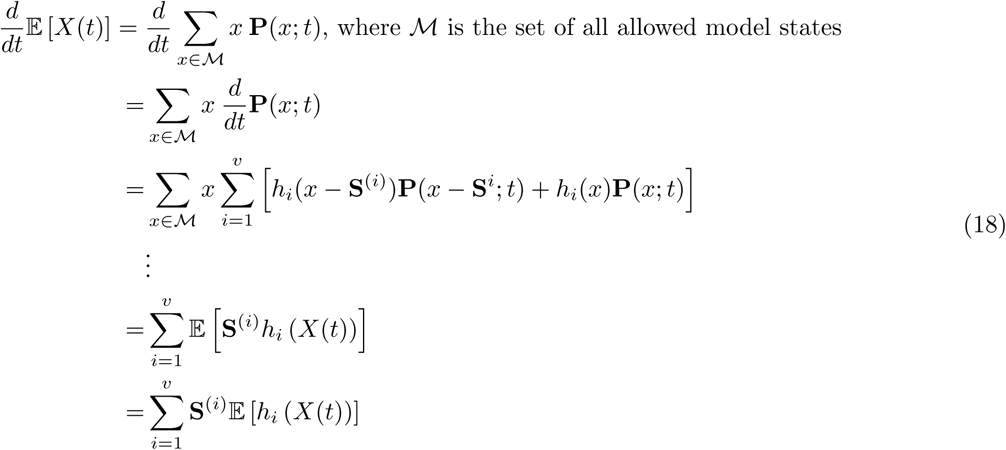

After making the substitution *y*(*t*) = 𝔼(*X(t)*), the above equation can be recognized as a matrix ODE giving the deterministic approximation of the system.

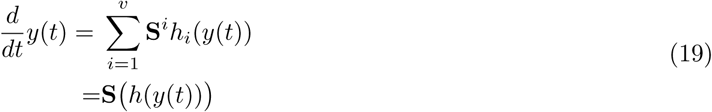

Substituting in our stoichiometry matrix **S** and SIS hazard functions, we derive the following matrix ODE (writing out the column vector *y*(*t*) explicitly in terms of our two state variables):

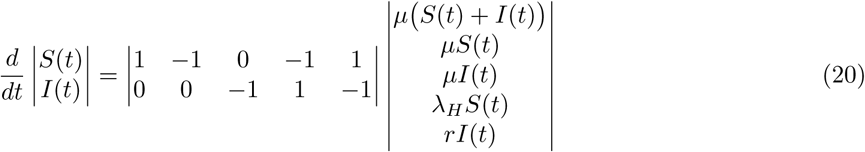

Separating the variables and completing the matrix vector multiplication leads to the familiar ODE form of the SIS model with demography. We defer the equilibrium solution until later, when we can solve for the mosquito equilibrium jointly.

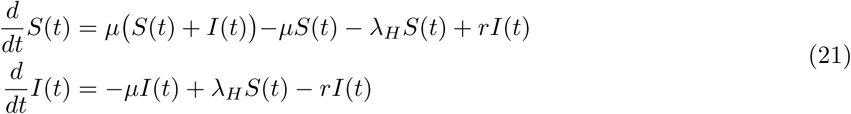

#### 2.2.2. Mean-field Approximation of Mosquito Stochastic Dynamics

Similar as we did for humans, we consider *λ*_*V*_ to be a constant and decouple the mosquito SEI dynamics from the human SIS dyanmics. A flow graph of the mosquito dynamics is shown in Figure S3, which shows the possible states a adult female mosquito may exist in during its life, with death (*D*) as an absorbing state. As described in the main text, we partition the exposure (extrinsic incubation period, EIP) into *n* compartments such that the overall dwell time is Erlang distributed. As before, because we are purely focused on the mosquito model, we drop the *V* superscript for state variables.

For a single adult female mosquito, the transition rates on the edges of the graph specify the hazard rates of leaving the current state; the aggregated process for a population of mosquitoes sums the individual hazards by the number of mosquitoes in that state.

**Table S1.**
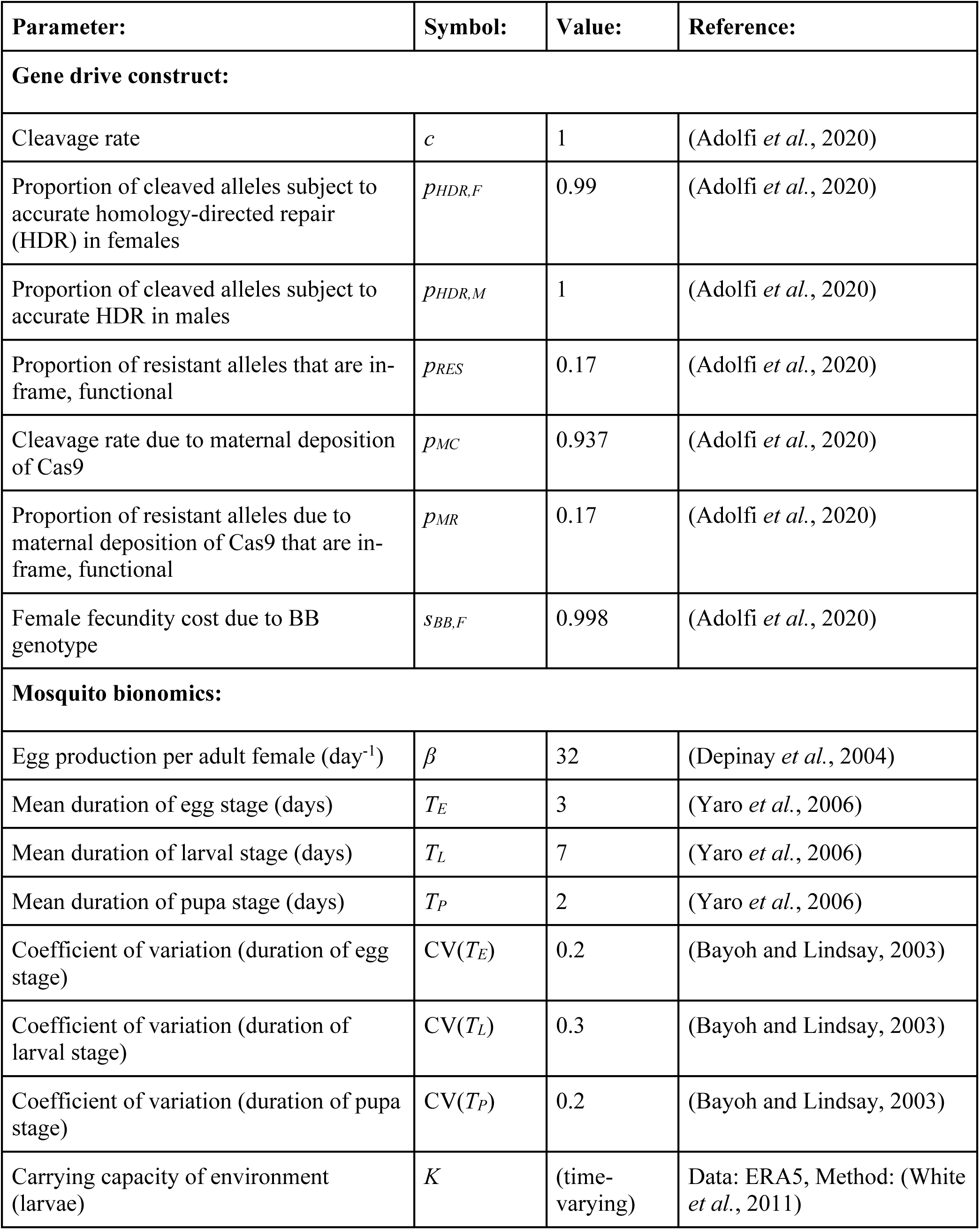
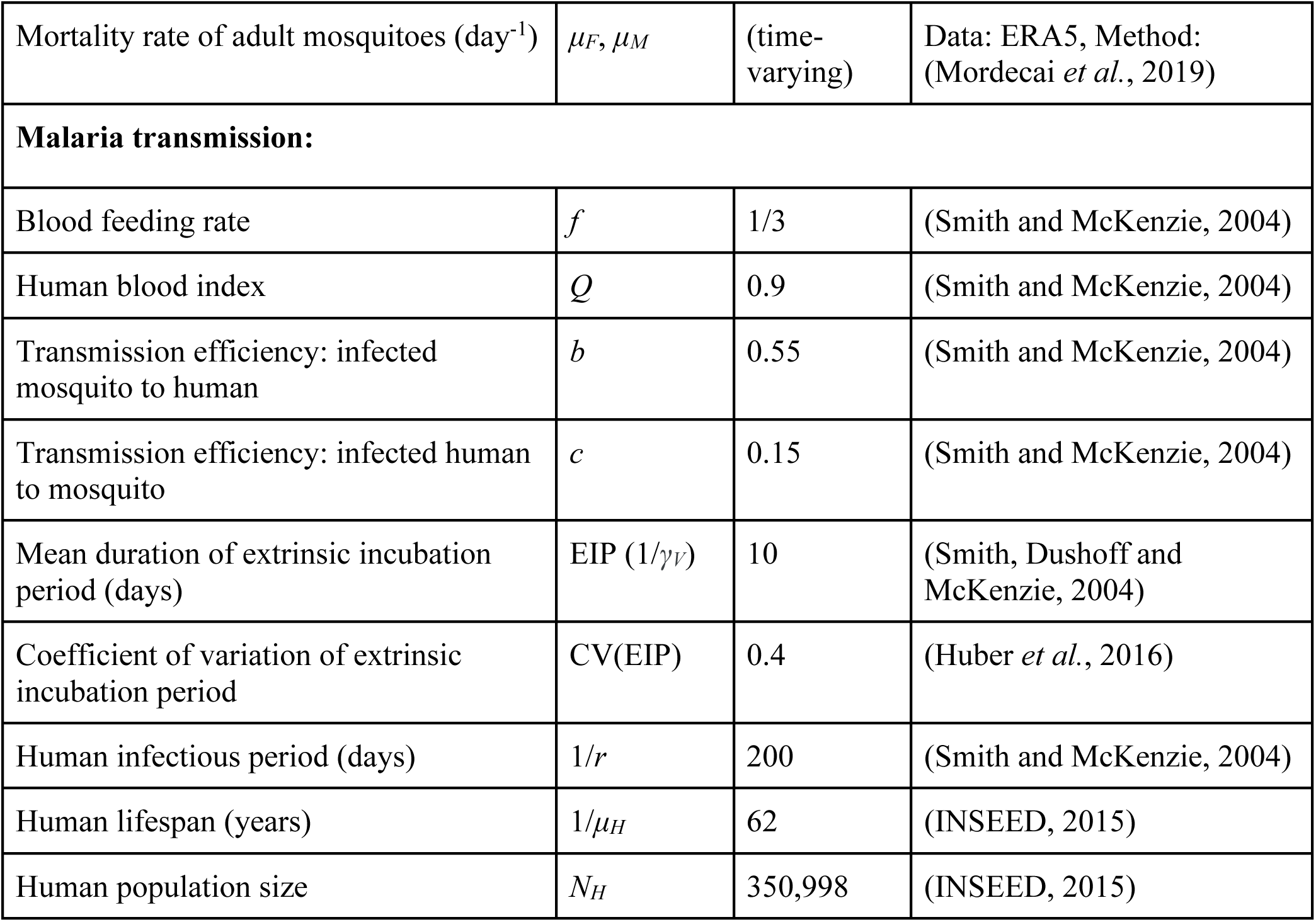
Model parameters describing the gene drive construct, mosquito bionomics and malaria epidemiology for simulations resembling releases on Grand Comore, Union of the Comoros.

**Figure S3:**
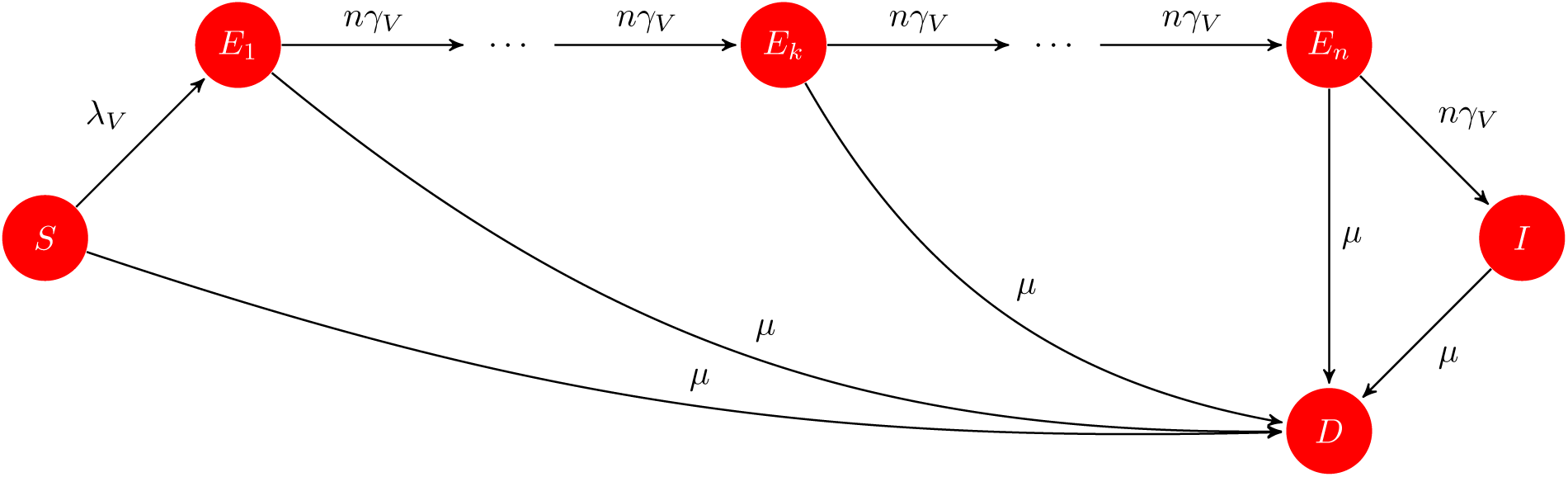
Susceptible-exposed-infected (SEI) mosquito infection dynamics

Upon emergence, the mosquito enters the susceptible *S* state, subject to force of infection *λ*_*V*_. Susceptible mosquitoes are also subject to a mortality rate, *µ*, which is constant across all compartments, leading to an exponentially distributed lifespan with mean 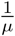. If the mosquito becomes infected, which occurs at rate *λ*_*V*_, it will advance through the extrinsic incubation period (EIP) prior to becoming infectious.

The EIP is broken into *n* bins, with transition from the *k*^th^ to *k* + 1^th^ occurring at a rate *nq*. This specification allows an Erlang (Gamma with integer shape parameter) distributed duration of EIP, with mean 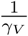 and variance 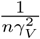. A mosquito survives the EIP with probability 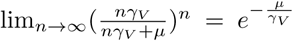 Conditional on survival, the proportion of mosquitoes that become infectious *t* days after becoming infected is distributed as Gamma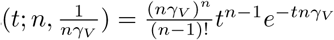 (using shape/scale parameterization).

Written in matrix form, the infinitesimal generator matrix for a single adult female mosquito, or a cohort emerging at the same time has the following form:

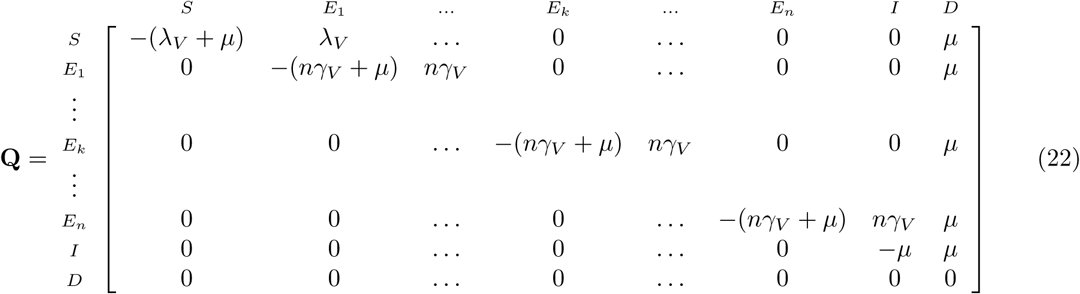

For the model of a single adult female mosquito, or a cohort that emerged at the same time, the infinitesimal generator (Equation 22), the KFE is:

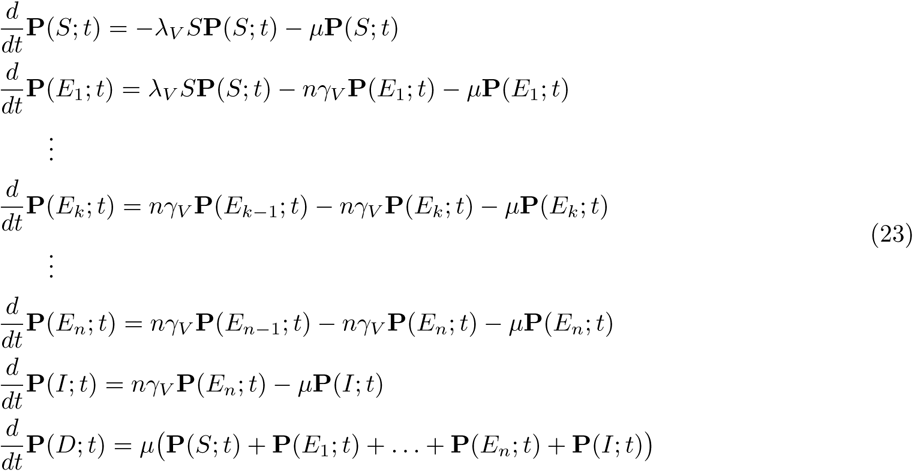

While Equation 23 describes how the probability distribution over states for a cohort of mosquitoes changes over time, to account for emergence (which we will need for the deterministic approximation), we let *E* give the rate at which females emerge into *S* from pupae. For brevity, **P**(…; *t*) appearing in the joint density function means that those elements of the random vector do not change.

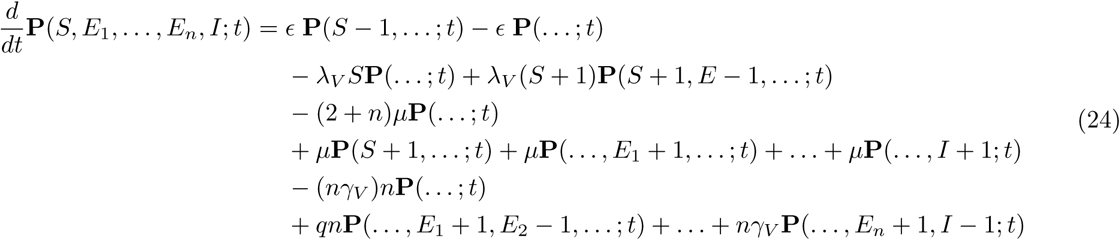

As we did for deriving the deterministic approximation of human infection dynamics, we write out the stoichiometry matrix **S**_*u×v*_ of dimensions (2 + *n*)×(2*n* + 4). Because each column in **S** describes an allowable jump in state space, the total number of terms in the KFEs should be equal to 2(2*n* + 4) = 4*n* + 8; checking this with the derivation in the previous section allows us to confirm that the equations are correct. Once we have **S**, the mean-field approximation follows the same method as in Section 2.2.1.

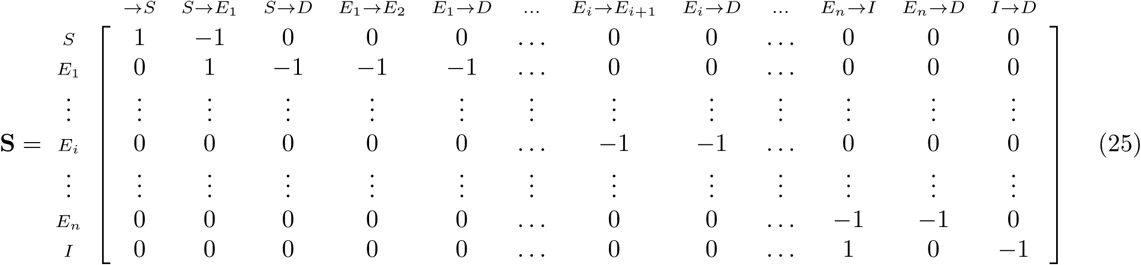

Using the stoichiometry and the KFEs we write down the approximating equations in matrix ODE form:

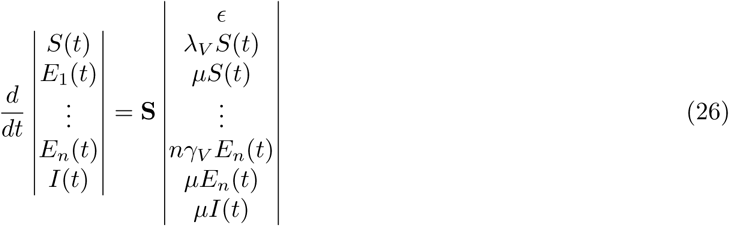

Because the emergence rate *ϵ* and the force of infection on mosquitoes *λ*_*V*_ are considered a constants, all jump terms are of zero or first order and the deterministic approximation will correctly approximate the mean behavior of the stochastic system. For clarity, we write the system of linear ODEs component-wise:

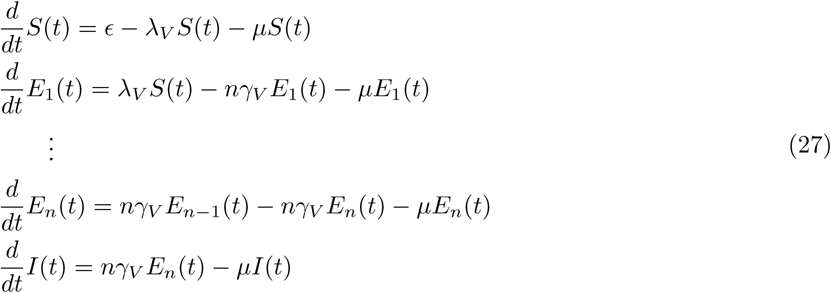

### 2.3 Quasi-stationary distribution for mosquito infection dynamics

In order to solve the coupled mosquito SEI - human SIS model at equilibrium, we need to be able to solve for the distribution of adult female mosquitoes across states (*S, E*_1_,…, *E*_*n*_, *I*). This is because, given an endemic equilibrium prevalence in humans, we can compute the number of infectious mosquitoes *I* required to sustain that prevalence of disease. From that, we can use the quasi-stationary solution of the CTMC model given in Equation 22 to compute the total adult female mosquito population and their distribution across stages, which can then be plugged into the life history equilibrium Equation 2 or Equation 4 to solve the entire model’s endemic equilibrium. We note that for the stochastic model, this is not a stationary distribution, but a quasi-stationary distribution (QSD), as there exist absorbing states in the model.

To compute the QSD, note death (*D*) is an absorbing state and the set of transient states is 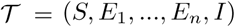. Then the random variable representing time to absorption (death) as a phase-type distribution. The QSD over 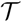 will arise by conditioning on survival. This distribution allows us to distribute mosquitoes across the transient states properly at equilibrium. We partition **Q** as (Bladt and Nielsen 2017; Buchholz, Kriege, and Felko 2014):

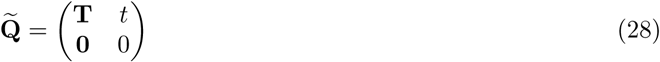

Here, 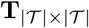 is a subintensity matrix of transition rates between transient states, and 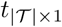 is a column vector of exit rates to the absorbing state. We denote the random variable following a continuous phase-type distribution describing time until absorption as *τ* ∼ PH(*π*, **T**) with density function *f*_*τ*_(*u*) = *πe*^T*u*^*t*, where 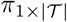 is a row vector specifying the initial distribution over transient states.

Let **U** = (− **T**)^*−*1^ be the matrix containing the means of random variables *uij* denoting time spent in state *j* starting from *i*, prior to absorption. We can use this matrix to define the QSD over 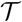, denoted as 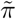. The j^th^ element of the quasi-stationary distribution is (Darroch and Seneta 1965; Darroch and Seneta 1967):

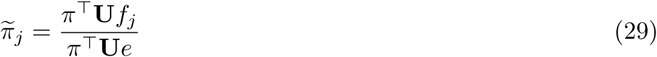

In Equation 29, *f*_*j*_ is a column vector with 1 in the j^th^ row and 0 elsewhere, and *e* is a column vector of 1’s. In our specific case, *π* places all mass on state *S* because the mosquito cannot emerge from the pupa stage already infected (there is no vertical transmission of pathogen), so the quasi-stationary distribution can be directly obtained from **U**.

#### 2.3.1 Coupled SEI-SIS Equilibrium Solutions

In order to solve for the mosquito population required to produce some equilibrium prevalence in humans, let *N*_*V*_ be the total adult female population, summing over infection states. Writing the SIS human dynamics (Equation 21) with the expanded form of *λ*_*H*_, we have:

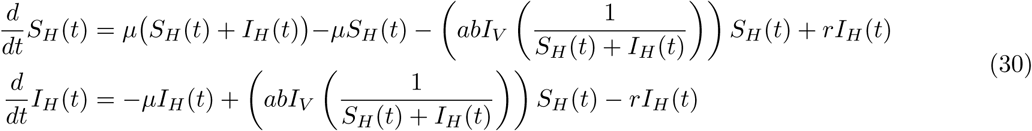

Because we consider both the human population size *N*_*H*_ = *S*_*H*_ + *I*_*H*_ and equilibrium prevalence 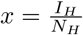 known constants, we can solve the model at equilibrium in terms of the number of infected mosquitoes, such that:

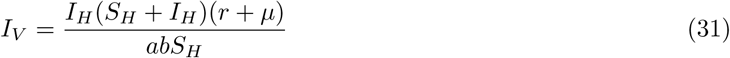

After we have solved for the number of infected mosquitoes *I*_*V*_, we can derive the total female mosquito population this implies:

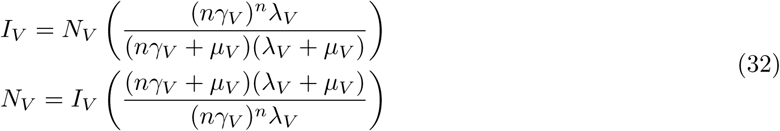

This total size *N*_*V*_ can be spread across the EIP stages via Equation 29. Given *N*_*V*_, we can calculate equilibrium solutions for the full lifecycle model by plugging in this number of adult females into either Equation 2 or 4.

### 2.4 SEIR Model

We also present results for an SEIR style model of human dynamics. An additional parameter, *γ*_*H*_, is the rate of progression from *E*_*H*_ → *I*_*H*_, that is, the inverse of the duration of latency in humans.

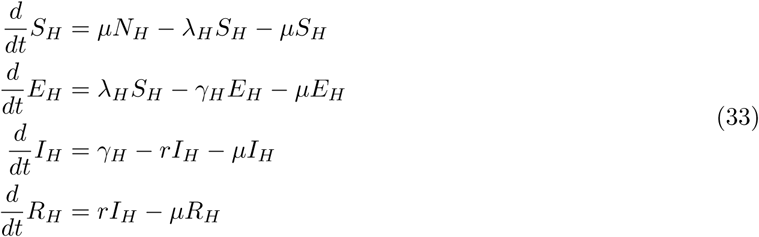

Where *N*_*H*_ is the total human population and the force of infection on humans follows the Ross-Macdonald form 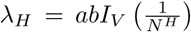. We consider that the total population size *N*_*H*_ and the number of infected and infectious humans *I*_*H*_ are known, allowing us to solve for *R*_*H*_ and *E*_*H*_ at equilibrium. It should be noted that for realistic parameter values, this model only has a non-trivial equilibrium when 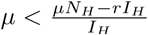, which results in extremely unrealistic (short) lifespans. The other two equilibrium points are the trivial disease free equilibrium when *λ*_*H*_ = 0 and the normal (*λ*_*H*_ > 0) case when *R*_*H*_(℞) → *N*_*H*_, that is, for realistic values of parameters, all surviving individuals will become infected subsequently recover. For that reason we do not explicitly calculate the endemic equilibrium. In general the SEIR human model should be used to evaluate the impact of gene drive interventions on one-off *epidemic* situations (eg; does releasing large numbers of modified mosquitoes 10 days after initial cases appear make a significant difference in final outcome), rather than for investigating *endemic* diseases, which require more complex models with waning immunity.

## 3 Stochastic Petri Net

Here we provide an introduction to the stochastic Petri net modeling formalism used in **MGDrivE 2**. Notation in this introduction is borrowed from (Wilkinson 2006).

### 3.1 Properties of SPN

SPN is a mathematical modeling language to describe discrete event systems, that is, a system which has a countable set of events (although only a finite number may enabled at any given time), each of which changes state in some way when it occurs. When events are assumed to happen after Exponentially-distributed intervals (alternatively, each event occurs at a constant, age-independent rate), the SPN is isomorphic to a CTMC, and can be extended to provide a modeling semantics for generalized semi-Markov processes (Glynn 1989). For practical application, a benefit of adopting the SPN modeling language is that model representation is separate from numerical simulation. This can allow both for highly efficient simulation, as the model can be represented via vectors and sparse matrices, and also utilization of model-agnostic simulation algorithms that take as input a generic SPN model and output sampled trajectories.

A Petri net is, formally, a bipartite graph, consisting of a set of *places*, 𝓟, and a set of *transitions*, 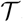. Directed edges, often called *arcs*, lead from places to transitions and from transitions to places. Arcs are allowed to have a positive integer weight. Therefore, if *u* = *|𝓟|* and 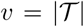, the set of arcs that connect places to transitions can be denoted by a non-negative integer matrix **Pre**_*v×u*_, and the set of arcs connecting transitions to places by **Post**_*v×u*_.

This bipartite graph so far defines the *structural* properties of the model. When translating conceptual models to the language of Petri nets, the places define the allowable state space of the model. However, in order to describe any particular state of the model, the Petri net must be given a *marking*, *M*, which is given by associating with each place a non-negative integer number of *tokens*, such that *M* ∈ *ℕ*_*u*_. Put more concretely, we can imagine taking some number of indistinguishable tokens and assigning each one to a place in the set 𝓟; the resulting vector in ℕ^*u*^ is a valid state of the model. In the language of CTMCs the marking *M* is referred to as the *state*, and we use the terms interchangeably.

Each transition 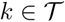 is allowed to change state when it occurs (fires). A transition *k* is *enabled* in some marking *M* and may fire when there are tokens on the place corresponding to each input arc (*k*th row of **Pre**) greater than or equal to the arc weight. When the transition fires, *M* is updated by removing tokens from M given by the set of input arcs and weight, that is, according to **Pre**. It then adds tokens in *M* according to its output arcs and weight in **Post**. This can be represented succinctly if we let **A** = **Post Pre**, then if *r*_*k*_ is a column vector of zeros with a one at place *k*, the state can be updated as *M* = *M* + **S***r*_*k*_, where **S** = **A**^T^. **S** has dimensions *u × v*, so it maps vectors in the space of events to vectors in the space of marking updates.

So far we have described a deterministic Petri net. However, associate with each transition a clock that tells us when *k* will fire, if it were the first of all enabled transitions to fire and let the process associated with *k* be a Poisson process *Y*_*k*_ with intensity *λ*_*k*_ which may depend on the current time *t*, and current marking *M* (*t*). Finally, if we let all enabled processes *Y*_*k*_ compete under a race condition by sampling the next firing time for each clock, *τ*_*k*_, such that *k*′ = arg min*k* {*τ*_*k*_}, then *k*′ is the event that fires. In that case the system time is updated to *t* = *t* + *τ*_*k′*_ and the state as *M* (*t*′) = *M* (*t*) + **S***r*_*k*′_. It can be rigorously proven that such a construction is a continuous-time Markov chain (Brémaud 1999).

An advantage of this construction of a Markov process rather than the more traditional presentation via the infinitesimal generator matrix is that processes with infinite state spaces can be compactly represented, because only a finite number of clock processes compete at any given time. In this way, infinite birth death processes, for example, can be succinctly represented graphically and simulated. Additionally, because most transitions only have a few input and output arcs, the matrices **Pre** and **Post**, which define the bipartite graph, can be sparse matrices, and the marking update step can use fast sparse matrix routines, enhancing computational efficiency.

### 3.2 SPN Architecture

We have developed algorithms to construct Petri nets for arbitrary genetic inheritance cubes (Sánchez C. et al. 2019), metapopulation structure, Erlang-distributed aquatic stages, infection dynamics, and human populations. Once built, and augmented with parameters for hazard functions in 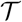, the resulting SPN model can be numerically evaluated via a variety of sampling algorithms. We describe the SPN architecture without considering epidemiological dynamics, as those are considered in a later section.

**MGDrivE 2** has been designed with consideration for computational efficiency. We store the matrices defining the SPN in sparse matrix format using the Matrix **R** package (Bates and Maechler 2019). In addition, when constructing 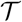, we check the input inheritance cube. If the viability mask 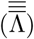 indicates a certain cross will never produce viable offspring, or if the probability of offspring for a cross is zero 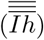, that transition is not instantiated in the SPN.

Generation of the set of places *P* for a single node is simple, and requires the user to pass the inheritance cube 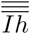, and the shape parameters for Erlang dwell distributions associated with egg, larval, and pupal aquatic stages to the function. The function returns the named set of places, along with an indexing data structure containing the indices of places stratified by life stage and genotype, both for easy debugging and the construction of arcs when the set of transitions is made. We note that SPN defines no particular order on the set 𝓟 but we “unroll” the set hierarchically into a vector first by node, life-stage, and genotype, for easier comprehension and testing.

After *𝓟* is constructed, we can construct transitions 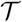, using *𝓟* as input, as well as the aforementioned shape parameters of aquatic dwell times and 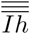. While not strictly necessary for SPN, we adopted several conventions here which simplify later generation of hazard functions. We first defined “classes”, K, of transitions, such that, for example, all transitions related to oviposition were grouped together. Each class *K* then is a proper subset of all transitions, 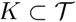, and 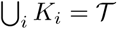. Each individual transition in a set *k*_*j*_ ∈ *K*_*i*_ has an associated **R** function which returns a data structure containing, at minimum, an index `vix`, indicating where in 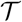, *k*_*j*_ can be found, a `label`, a character string giving the name of the transition, `s` and `as_w`, indices of input arcs (the places they originate at), and weights respectively, `o` and `o_w`, the same for output arcs from this transition back to places, and `class`, giving the name of the class *K*_*i*_ as a character string this transition belongs to. To make 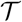, we iterate through classes *K*_*i*_, and *k*_*j*_ within classes, storing each transition’s packet of information in the vector 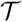. Because each transition “knows” its input and output arcs, as well as their weights, adding new classes of transitions is simple, as a single function merely needs to be written that takes in places and perhaps genetic information, and returns this minimal packet of information.

Once 𝓟 and 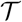 are constructed, the SPN is formally constructed. However for computation, we prefer to store a more compact representation of the net. It is at this point we build the sparse matrices **Pre** and **Post**. To do so, we simply allocate two *v × u* sparse integer matrices, then iterate through 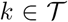. We use the information packet described earlier, specifically `s, s_w` and `o, o_w`, to fill in the non-zero entries of **Pre** and **Post** respectively. Because we have already induced ordering on 𝓟 and 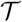, the matrices have the right sorting of rows and columns.

### 3.3 Hazard Functions

To build a CTMC model from the Petri net (SPN), each transition must have a hazard function *λ*_*k*_ (*t, M* (*t*)). In the code implementation of **MGDrivE 2**, each hazard first checks if that transition is enabled, if not it immediately returns zero and else computes the hazard rate. We allow *λ*_*k*_ to be a function of time for simulation of inhomogeneous processes. Because in this manuscript we consider only Markovian systems, hazards only depend on the current system state *M* (*t*) and time *t*. Unless otherwise noted, we use *λ*_*k*_ to generically denote the joint hazard and enabling function for process *Y*_*k*_.

This use of CTMC is known as a Markov population process, and has been used to model stochastic population models for some time now (Kingman 1969). However, it can be useful to include exogeneous stochastic processes into the model, which may affect hazard rates. These processes could represent, for example, environmental processes such as temperature or rainfall. Knowledge of these processes would be necessary to evaluate the hazard functions. Consider the situation in which process *Y*_*k*_ is affected by environmental stochasticity represented by *z* so that the hazard is *λ*_*k*_ (*t, M* (*t*), *z*); for concreteness, consider *z* to be temperature and *k* to be larval mortality. We must also consider a specific function of interest (*f*) to be computed from trajectories of **MGDrivE 2**, which we would like to estimate via Monte Carlo, to average over uncertainty in *z*. Again for concreteness, we could consider functions like time required for a specific gene to fixate, or time required for pathogen extinction in a specific node. To propagate uncertainty from arbitrary exogenous processes, we simply draw many samples from *z* and then run Monte Carlo simulation of **MGDrivE 2** on each realized exogenous trajectory; we can imagine an “outer” loop sampling a trajectory 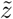 from *z* and an “inner” loop computing Monte Carlo estimates of *f*, conditioning on 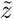 as deterministic input to 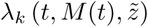. Because *z* is by definition an exogeneous source of stochastic variation, the probability factorizes such that this method properly propagates uncertainty into our estimation of *f*. For specific functions, more efficient methods than naive Monte Carlo may exist, and we refer to the panoply of variance-reduction methods covered in (Bratley, Fox, and Schrage 2011).

In **MGDrivE 2**, once the Petri net 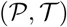 is constructed (and we have parameters *θ* and inheritance cube 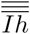), we can construct the *v*-dimensional column vector of hazard functions Λ. Specifically, we store the individual *λ*_*k*_ functions as function closures within the vector Λ, as functions are first class objects in **R**. We note that this can be easily adapted to other programming languages, for example in C++98, functors could be used in lieu of closures, and in C++11/14, lambda functions could achieve the same effect (Meyers 2014). Each closure stores only the elements of *θ* and 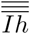 necessary for computation of the hazard. We note that the function closure based storage of hazard functions means that it is easy to include additional computational state for specialized algorithms or more complex processes, such as enabling times or integrated hazards.

So far we have described what the fully constructed object Λ is, but not yet how we implemented it in code, which we do now. Much like the construction of 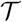, we rely heavily on our assignment of transitions into classes; after allocating memory for a *v*-length vector, we begin iterating through 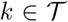. First, we scan the class of the transition; this tells us the appropriate function factory to call that will return *λ*_*k*_, the function closure that computes the hazard. We pass the packet *k* to the function factory, along with parameters *θ*. The packet provides all the necessary information to set up the enabling rule, and the function factory pulls out the necessary components of *θ* to compute that specific hazard, which are stored in the function closure. All returned hazard functions *λ*_*k*_ accept only two arguments, *t* time, and *M*, the state (marking).

We allow the option for users to select if the vector of hazard functions shall be “exact” or “approximate”. Exact hazards are required for both sampling algorithms that simulate integer numbers of tokens, in which case enabling rules make sense. In this case evaluation of *λ*_*k*_ proceeds as described above. If however a continuous-state approximation is desired, either from a deterministic interpretation of the hazard functions as rate functions for mean-field approximation, or from a drift-diffusion stochastic differential equation, we ignore the check for sufficient tokens on input arcs, as the model no longer has an integer state space.

### 3.4 Numerical Simulation

One key feature of the SPN representation of **MGDrivE 2** is the convenient decoupling of model specification and sampling method, allowing model-independent development of fast algorithms. This lets us benefit from extensive work into optimized stochastic sampling algorithms from the chemical kinetics and physical simulation communities, many of which can be used nearly “off the shelf” with a place/transition model representation. We refer to the encyclopedic book by (Marchetti, Priami, and Thanh 2017) as one of many resources for fast simulation routines.

Currently we do not support exact simulation of inhomogeneous processes, although approximate simulation is best done via the Poisson time-step method, where inhomogeneous terms are discretized to a piecewise constant step function with the same ∆*t* as used in the approximate time-step. Exact simulation of inhomogeneous processes is difficult, although an algorithm based on random time change was presented by (Anderson 2007), and (Thanh et al. 2018) investigate exact and approximate rejection-based methods. We leave the incorporation of these or similar sampling methods into the **MGDrivE 2** framework for future development.

We provide example code to numerically integrate deterministic trajectories, based on the deSolve **R** package of ODE solvers (Soetaert, Petzoldt, and Setzer 2010), using a mean-field approximation to the stochastic system (Bortolussi et al. 2013). We also provide several stochastic samplers for both exact and approximate trajectories, inspired by the smfsb **R** package (Wilkinson 2006).

In certain situations, when populations are large (that is, no places have a small number of tokens) and hazard functions are close to linear (guaranteed when using mass-action forms), it may be the case that stochastic fluctuations can be safely neglected. (Kurtz 1970) made rigorous the conditions under which CTMCs may converge to ODEs. Such an approach essentially simplifies to considering the hazard functions as rate functions, and the state *M* (*t*) as a continuous quantity (motivating the ability for the user to select generation of “approximate” Λ) (Gillespie 2007). Even when stochastic fluctuations are non-negligible such that deterministic approximation is not valid, it may be useful to provide the option for deterministic integration of the SPN model for quick visualization of transient behavior, or for sensitivity analysis.

At present we only offer Gillespie’s “direct-method” to sample statistically exact trajectories, a well known method to sample from stochastic models (Gillespie 1977). Being an exact sampler, it samples integer-valued trajectories, and thus uses exact hazards and enabling functions. Briefly, the method works via a simple update step where first the vector of hazard functions is evaluated, *h(t)* = Λ (*t,M(t)*). The Markov transition kernel to the next state can be factored such that the sampler first samples the random variable *τ* describing when the jump occurs, relative to *t, τ* ~ (Σ_*k*_*h*_*k*_(*t*)). Next it samples *which* process caused the jump, and updates the system accordingly, that is, it selects the process causing the jump, *k*′ with probability 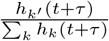. Then update the marking according to matrix equation *M*′(*t + τ*) = *M*(*t*)+**S***r*_*k*′_. In general, large populations of mosquitoes and/or large numbers of nodes would render exact simulation practically impossible, if, for example many tens of thousands of individual events needed to fire each day, each requiring a system update and resampling of random variables for each event, both of which are computationally expensive tasks.

In addition, there are two approximate stochastic sampling algorithms that have been implemented for use in **MGDrivE 2**, and we anticipate future algorithmic development focusing on implementation of improved approximate samplers. Th
e first of these is a simple fixed size tau-leaping method, the Poisson time-step (PTS), reviewed in (Wilkinson 2006) and first introduced by (Gillespie 2001). The basic concept behind the PTS algorithm is that if none of the hazard functions change significantly over a small time step, say [*t, t* + ∆*t*), then one can approximate the state change by sampling a Poisson distributed random variable for enabled each 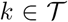, such that the elements of the *r* vector indicating how many times each event fired are each independent Poisson random variates with rate parameter *λ*_*k*_ (*t, M* (*t*)) ∆*t*. Then the matrix update can be preformed with those sampled Poisson variates in the vector *r*, system time updated, and another iteration preformed. The extent to which the assumption that hazards do not change significantly over the interval determines the quality of the approximation. The original tau-leaping algorithm has spawned many variations on the theme, including some with strong probabilistic guarantees of approximation quality (Anderson 2008), which may be incorporated for **MGDrivE 2**.

The second approximate stochastic sampling algorithm is based on a continuous state stochastic differential equation (SDE), known as the diffusion approximation. We offer a brief heuristic explanation of the algorithm but a detailed derivation can be found in (Gillespie 2005). If one starts with the integer valued Markov jump process, it will have a set of differential equations known as Kolmogorov’s forward equations (KFE), sometimes referred to as (chemical) Master equation. The KFE gives the complete time-evolution of the probability mass function across allowable system states, and is usually intractable. It is possible (see (Toral and Colet 2014) for a brief overview) to generate a partial differential equation (PDE) second-order approximation to the KFE known as the Fokker-Planck equation, which approximates the probability mass function with a probability density function evolving according to first order drift and second order diffusion coefficients. Considering the marking *M* as a continuous state, the Fokker-Planck equation can be interpreted as a *u*-dimensional SDE driven by independent Wiener processes. While advanced techniques for simulation of SDEs exist (Särkkä and Solin 2019), we implement the simple Euler-Maruyama method in **MGDrivE 2**.

